# Cue-Invariant Geometric Structure of the Population Codes in Macaque V1 and V2

**DOI:** 10.1101/2023.12.05.570110

**Authors:** Corentin Massot, Xiaoqi Zhang, Zitong Wang, Harold Rockwell, George Papandreou, Alan Yuille, Tai-Sing Lee

**Author notes:** Equal contribution (Corentin Massot, Xiaoqi Zhang, Zitong Wang). George Papandreou contributed to this project while employed at UCLA. He is currently working at Snap Inc., London, United Kingdom.

## Abstract

Our ability to recognize objects and scenes, whether they appear in photographs, cartoons, or simple line drawings, is striking. Studies have shown that infants, members of isolated Stone Age tribes, and non-human primates can readily identify objects from line drawings. These findings suggest that the brain may inherently generate neural representations that align across different rendering cues, enabling abstraction. To test this hypothesis, we investigated the representational invariance of complex patterns of surface boundaries found in natural scenes. We tested whether individual neurons in V1 and V2 of the macaque monkey responded similarly to the presentation of these patterns across different renderings (i.e., as contours, luminance-defined patches, and segments of natural images). We found that individual neurons exhibit some degree of tuning invariance, stronger in V1 than in V2. At the population level, as a means to assess cue-invariant abstract representation, we measured decoding accuracy across cues (‘cue-transfer’ decoding). We found that this decoding is greatly enhanced when a geometric transformation (Procrustes Transformation) is first performed to align the population activities across cues. It is also effective when applied to different populations of neurons within or across visual areas. These results were compared with populations of artificial neurons from models of the ventral visual streams, further indicating that cue-invariance stabilizes with population size. In summary, we found that while individual neurons exhibit some cue-invariance properties, the stability of the population geometry emerges as a more robust candidate for supporting a cue-invariant representation of visual information in the early visual areas.

**SIGNIFICANT STATEMENT:** How can we easily recognize objects and scenes in a wide range of renderings, such as photographs, cartoons, or line drawings? One possibility is that our visual system processes information using an invariant representation. To investigate this hypothesis, we designed a stimulus set made of boundary patterns extracted from natural scenes, and displayed using three distinct renderings. We found that, while the tuning preference of individual V1 and V2 neurons displayed some correlation across renderings, a more robust invariant representation could be achieved when analyzing neural population geometry. Overall, we found that a cue-invariant representation of visual elements in the early visual areas may rest primarily on the geometry of the population responses, rather than individual neurons’ tuning characteristics.

---

We can perceive objects and scenes whether they are in their natural environment or when depicted in photographs. We can perceive these same objects and scenes just as effortlessly when they are depicted using artificial rendering such as line drawings, sketches, and cartoons. This visual ability has been observed in human babies (Yonas and Arterberry, 1994), members of Stone Age tribes with no experience of image-based communication tools (Sayim and Cavanagh, 2011), and in animals, such as chimpanzees (Tanaka, 2007). This leads to the hypothesis that the visual system builds an invariant representation of objects through shapes defined by object surface boundaries, irrespective of their physical support, a process considered part of a core cognitive ability to represent elements of the visual environment (Spelke, 2022).

There is, however, a tension between abstract representation and the need for the visual system to maintain the sensitivity to the physical attributes of the stimuli, resulting in a differential trade-off between selectivity and invariance in the successive cortical areas along the ventral visual stream (Liu et al., 2025; Quiroga et al., 2005; Rust and DiCarlo, 2010). While selectivity has often been associated with tuning properties of visual neurons, many works have focused on elucidating what type of invariance the visual system was able to achieve and how cortical circuits could implement it. Neurophysiological evidence suggests that surface and object boundary information is computed and represented explicitly in the early visual cortex, V1 and V2. V1 has been shown to encode oriented segments (Heitger et al., 1992; Hubel and Wiesel, 1968) but also more complex shapes (Dobbins et al., 1987; McManus et al., 2011; Ponce et al., 2017; Tang et al., 2018; Victor et al., 2006; Vinje and Gallant, 2000). Recurrent circuits and contextual modulation in V1 have been extensively studied in the context of contour completion and association field (Angelucci et al., 2017; Fitzpatrick, 1996; Gilbert et al., 2000). More recent evidence suggests that V1 neurons encode even more complex patterns associated with boundary concepts such as corners, curves, and rings (Tang et al., 2018). One well-known invariant property of V1 neurons is the contrast invariance of their orientation tuning. This invariant property was originally described by Hubel and Wiesel in their seminal work (Hubel and Wiesel, 1968), and was followed by numerous studies that aimed at understanding the neural circuitry in V1 complex cells that underpins this property (Alitto and Usrey, 2004; Shams and von der Malsburg, 2002; Skottun et al., 1987; Troyer et al., 1998). V2 has also been shown to encode complex shape elements and composition of orientation features (Anzai et al., 2007; Hegdé and Van Essen, 2000, 2007; Roe and Ts’o, 2015) and be sensitive to texture patterns (Freeman et al., 2013; Laskar et al., 2020; Ziemba et al., 2016, 2018, 2019). These studies have demonstrated the capacity of the visual system to achieve some degree of rotation and translation in order to cope with the diversity of the visual information while still achieving recognition (Tacchetti et al., 2018). To date, some debate remains as to whether invariant processing is achieved through extreme sparse coding of individual neurons or population coding strategies (Kanwisher and Yovel, 2006; Lieber et al., 2025; Liu et al., 2025; Oleskiw et al., 2025; Quiroga et al., 2005; Vinje and Gallant, 2000).

Still, questions remain about the capacity of early visual neurons to create an abstract representation of surface boundary concepts. Do early visual cortical neurons show representational invariance of surface boundary across renderings? How can these neurons achieve such invariance? To address these questions, we investigated the extent to which V1 and V2 neurons encode an abstract invariant representation of a set of surface boundaries extracted from natural scene images, at both the individual-neuron and population levels. Our study extends classical cue-invariant studies in four new directions. First, we created a dataset by extracting ‘surface boundary’ primitives from 2D natural scenes (Chen et al., 2013). We chose transitions between different surfaces or objects because these visual elements have varying 2D shapes that reflect the natural statistics of the visual environment. A key advantage of this approach is that such 2D surface boundaries could be visually rendered in three different ways: edge segments, luminance-defined boundaries, and texture-defined boundaries. We measured cue-invariant representation in V1 and V2 based on the correlation between the tuning curves of individual neurons responding to these boundary patterns in the different renderings. Second, we examined the cue-invariant representation in V1 and V2 at the population level. We developed a cue-transfer decoding approach to assess cue-invariance in the geometric structure of population responses. To do so, we applied a linear classifier to classify each boundary pattern based solely on the neural population activity (i.e., we trained a ‘decoder’). If the classification accuracy is sufficiently high, it is assumed that the information content of the stimuli can be subsequently accessed (i.e., ‘decoded’) by a downstream cortical region. In our cue-transfer decoding approach, we trained this decoder on population responses to one rendering and tested its ability to decode the responses to the same surface boundaries presented in another rendering. Third, we found that the population code’s geometry is typically rotated, scaled, and translated across different renderings. Applying a Procrustes Transformation (PT), which improves the geometrical alignment of the population codes, reveals a stable underlying geometry that is preserved not only across renderings but also across different populations of neurons, both within and across visual areas. Fourth, we compared neuronal responses with those of units in artificial neural networks, using this comparison to estimate the population size required to achieve a robust, cue-invariant representation. Taken together, our results show that cue-invariant representation is weak in individual neuronal tunings of V1 and V2, as the population codes themselves rotate, scale, and translate across renderings. Despite this variability, the geometry of the encoding manifold is highly stable. We therefore suggest that the abstract cue-invariant representation of boundary information, which underlies our ability to perceive objects across different modalities, is encoded in the stable geometry of population responses within the early visual cortex. We conclude by discussing the relevance of our work to the broader fields of population geometry analysis and the brain’s abstract representation of visual information.

## Results

We recorded the responses of V1 and V2 neurons to the presentation of surface boundary stimuli in three renderings. The stimuli consisted of patches containing surface boundaries extracted from 2D natural scenes of the Berkeley segmentation database (Chen et al., 2013; Martin et al., 2001) (see Methods, Fig.1.a). The boundary elements within each patch had varying shape complexity, such as oriented lines, curves, and junctions, reflecting the natural statistics of the visual environment (Elder, 2015; Elder et al., 2018; Geisler and Perry, 2009). After applying edge segmentation, the patches were grouped into 50 clusters based on a pairwise shape-distance metric of the segmented contours. The 50 stimuli at the cluster centers were selected to create the ‘Edge Contours’ (EC) rendering dataset (Fig.1.b left). This rendering is reminiscent of a sketch or cartoon representation of object contours. Each ‘Edge Contours’ stimulus had a corresponding ‘Example’ stimulus, which was the original patch extracted from the natural scene. Together, these 50 ‘Example’ stimuli formed the ‘EX’ rendering dataset (Fig.1.b middle). This rendering preserved all the texture statistics and visual features of the original natural scene. Finally, principal component analysis was applied to each of the 50 patch clusters. The first component of each cluster was selected, yielding 50 ‘Appearance Contours’ stimuli that formed the ‘AC’ rendering dataset. (Fig.1.b right). This rendering captured the mean luminance and contrast information surrounding the boundary element in the original natural scene. Animals were trained to perform a visual fixation task during which a stimulus from the dataset was displayed on a screen and V1 or V2 neural activity was recorded (Fig.1.c). To compare the cue-invariance properties of V1 and V2, we set the size of the stimulus to the average size of the receptive fields in each region (1^°^ for V1 and 1.5^°^ for V2, Fig.1.c). In total, we recorded the activity of 157 V1 neurons (48 in monkey FR and 109 in monkey KO) and 258 V2 neurons (112 in monkey FR and 146 in monkey KO) in two non-human primates using multi-contact linear electrodes (see Methods).

**Figure 1:**
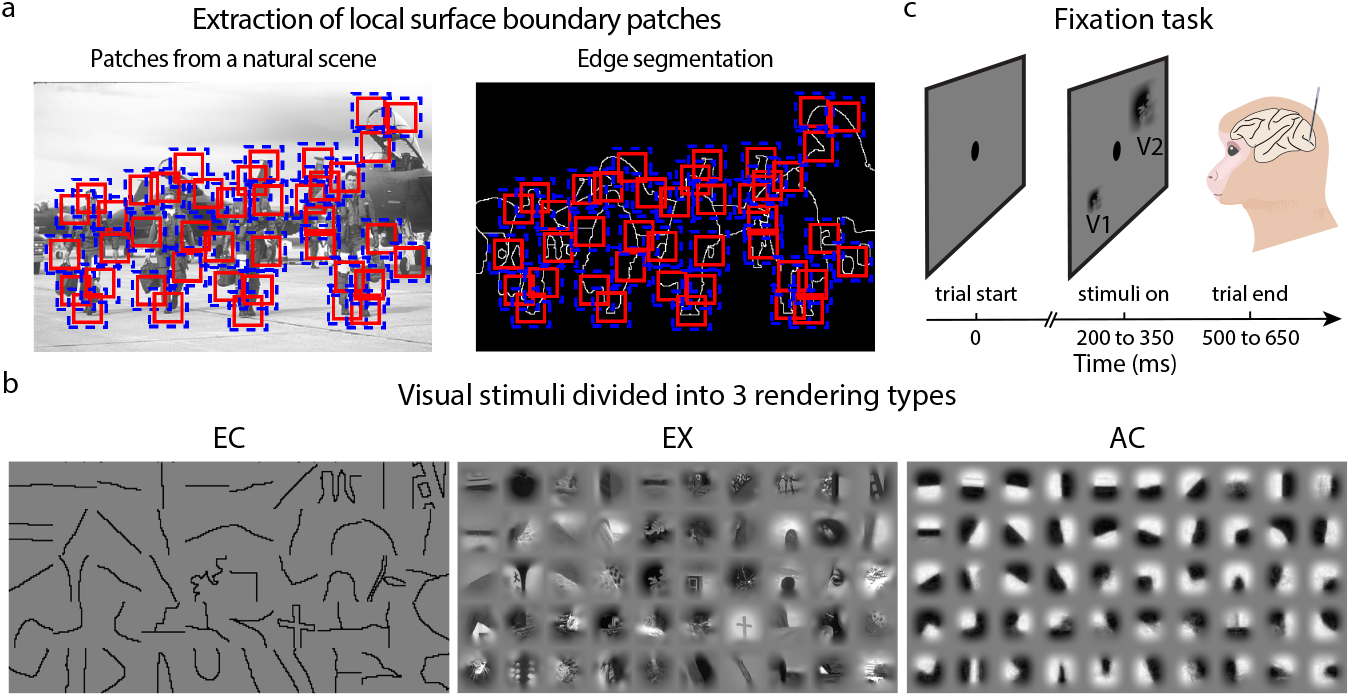
Visual stimuli and experimental setup. (a) Extraction of local surface boundary patches from a natural scene. Left: Example patches containing original image information are extracted from a natural scene. Right: The same patches are shown after edge segmentation has been applied to isolate the boundary elements within each patch. (b) The visual stimuli divided into three rendering types: ‘Edge Contours’ (EC, left), ‘Examples’ (EX, middle), and ‘Appearance Contours’ (AC, right). (c) Experimental setup: During the task, a laminar probe was acutely inserted to record population activity from either V1 or V2. Each trial began with the appearance of a central fixation dot. After a variable delay, a single stimulus was displayed, with its size differing between V1 and V2 recordings. The animal was required to maintain fixation on the dot until the stimulus disappeared for the trial to be labeled correct and to receive a liquid reward.

### Tuning Correlations between Stimulus Renderings of Individual V1 and V2 Neurons

First, we examined cue-invariance properties of individual neurons in V1 and V2. Fig.2.a shows the three sets of stimuli, each rank-ordered according to the trial-averaged firing rate of an example V1 neuron recorded in monkey KO. A visual inspection of the first row in all three panels reveals that this neuron displays increased responses to the same surface boundary stimuli across the different renderings. When plotting the rank-ordered responses for each rendering, we obtain three ‘tuning curves’ (Fig.2.b). As these tuning curves do not overlap, a comparison between renderings based solely on the trial-averaged firing rates does not capture any invariance property. Instead, we analyzed the preservation of the relative stimulus preferences between renderings, which represents a cue-invariance property of individual neurons.

**Figure 2:**
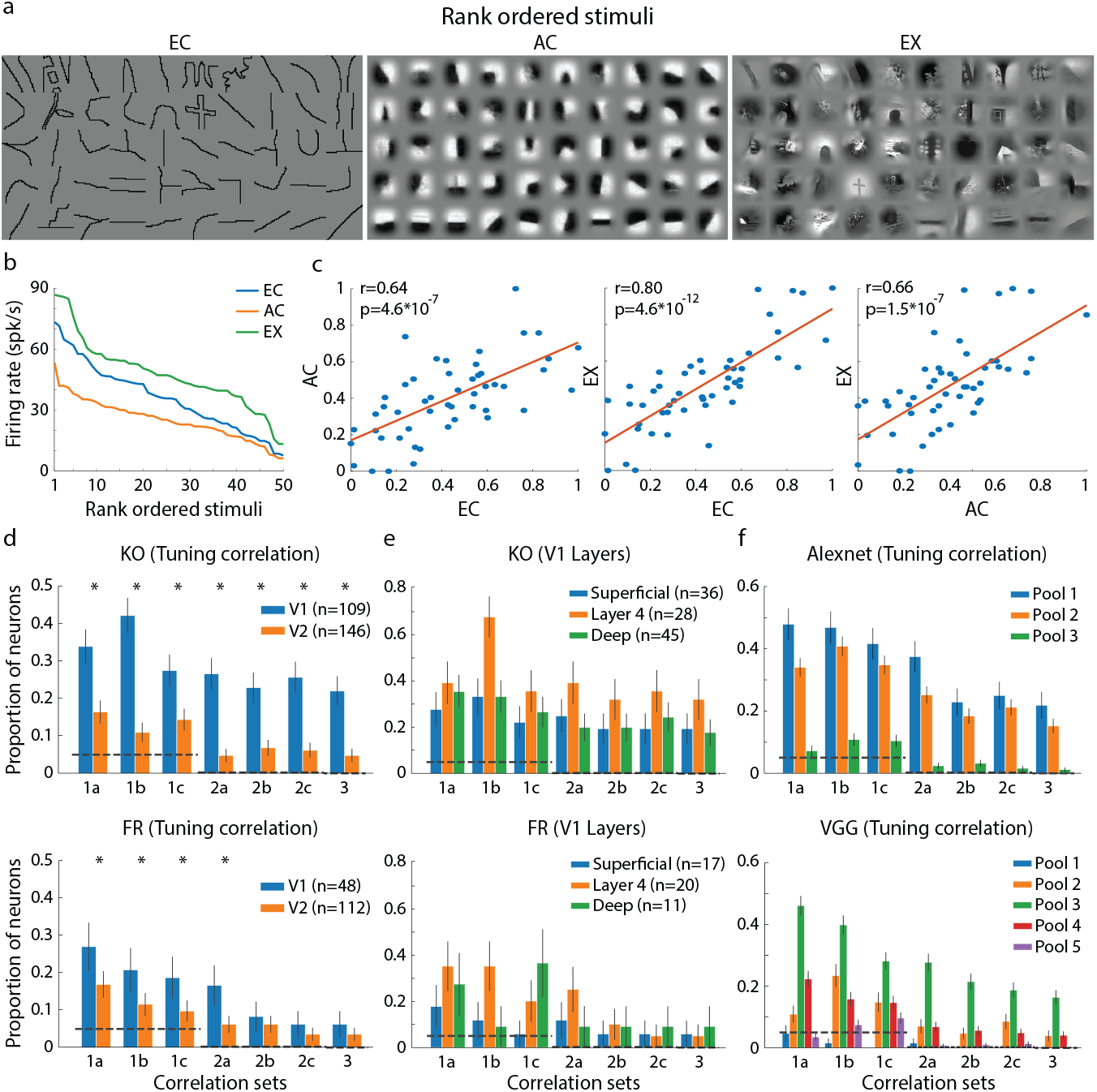
Tuning correlation of individual neurons in V1 and V2. (a-c) Tuning correlation analysis of an example V1 neuron in monkey KO. (a) Stimuli for each rendering, rank-ordered by the neuron’s trial-averaged firing rate. (b) Tuning curves of the example neuron for each rendering. (c) From left to right, scatter plots showing the neuron’s responses to all stimuli for the 3 rendering pairs (EC-AC, EC-EX, AC-EX). The red line indicates the correlation between a pair of tuning curves (i.e., tuning correlation), with the Pearson correlation and p-value shown in the insets. (d) Proportion of V1 (blue) and V2 (orange) neurons with significant tuning correlations, shown for different correlation sets in monkey KO (top) and FR (bottom). Correlation sets are: EC-AC (1a), EC-EX (1b), AC-EX (1c), EC-AC and EC-EX (2a), EC-AC and AC-EX (2b), EC-EX and AC-EX (2c), EC-AC and AC-EX and EC-EX (3). We also group these sets by the number of pairs they contain: one-pair (1a, 1b, 1c), two-pair (2a, 2b, 2c), and three-pair (3). Error bars indicate SSE (the standard error of sample proportion) confidence intervals. Horizontal dashed lines indicate the chance level for correlation sets with one-(*p* = 0.05), two-(*p* = 2.5 × 10^−3^), and three-pair (*p* = 1.25 × 10^−4^) rendering sets. Asterisks (*) denote a significant difference between the V1 and V2 proportions (using CPP, Comparison of Population Proportion). The legend indicates the number of neurons in each area. (e) Tuning correlation results for V1 neurons grouped by cortical layer: Superficial (blue), Layer 4 (orange), and Deep (green). Panel details are as in (d). (f) Tuning correlation results for artificial units from 3 pooling layers of AlexNet (top) and 5 pooling layers of VGG (bottom). Panel details are as in (d).

**Figure 3:**
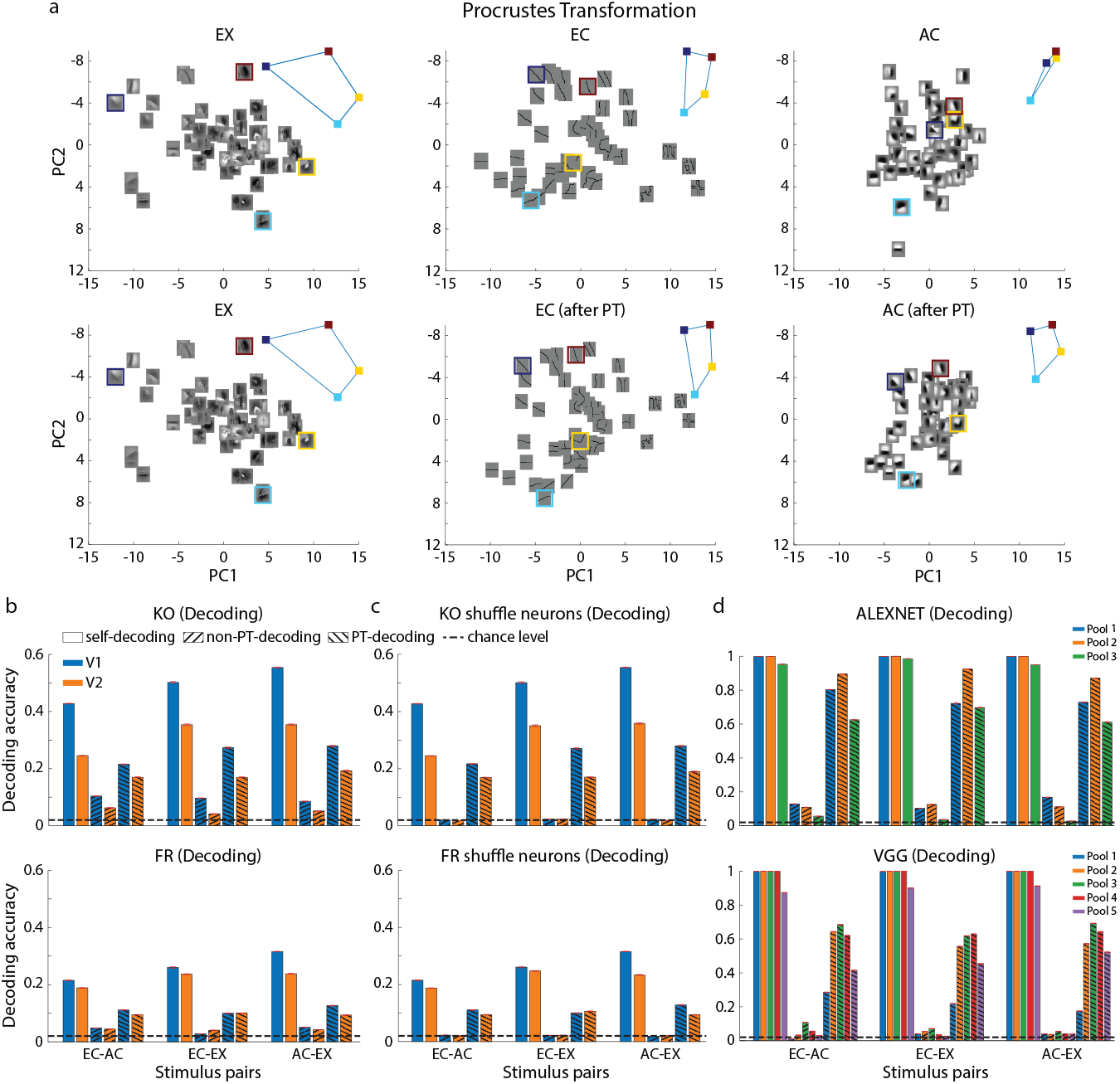
Cue-transfer population decoding and Procrustes Transformation. (a) Example of the V1 population (monkey KO) response geometry for all stimuli in each of the three renderings, projected onto the first two principal components (PCs). The top row shows the raw geometries. Bottom row shows the same geometries after the Procrustes Transformation is used to align the EC (middle) and AC (right) renderings to the EX rendering (target). Insets highlight four example stimuli (colored borders) to show their improved alignment after the transformation. (b) Cue-transfer population decoding accuracy for V1 (blue) and V2 (orange) in monkeys KO (top) and FR (bottom). The x-axis shows the rendering pair used for transfer decoding. For each pair, three conditions are shown: self-decoding (plain bars, an upper bound), non-PT-decoding (forward-slash hatching, transfer decoding without PT), and PT-decoding (backslash hatching, transfer decoding with PT). The horizontal dashed line indicates the 2% chance level. Red error bars show the SEM (the standard error of the mean). (c) Cue-transfer decoding after shuffling the neuron-by-neuron correspondence. Panel details are as in (b). (d) Decoding performance for populations of artificial units from AlexNet (top) and VGG (bottom). Panel details are as in (b).

**Figure 4:**
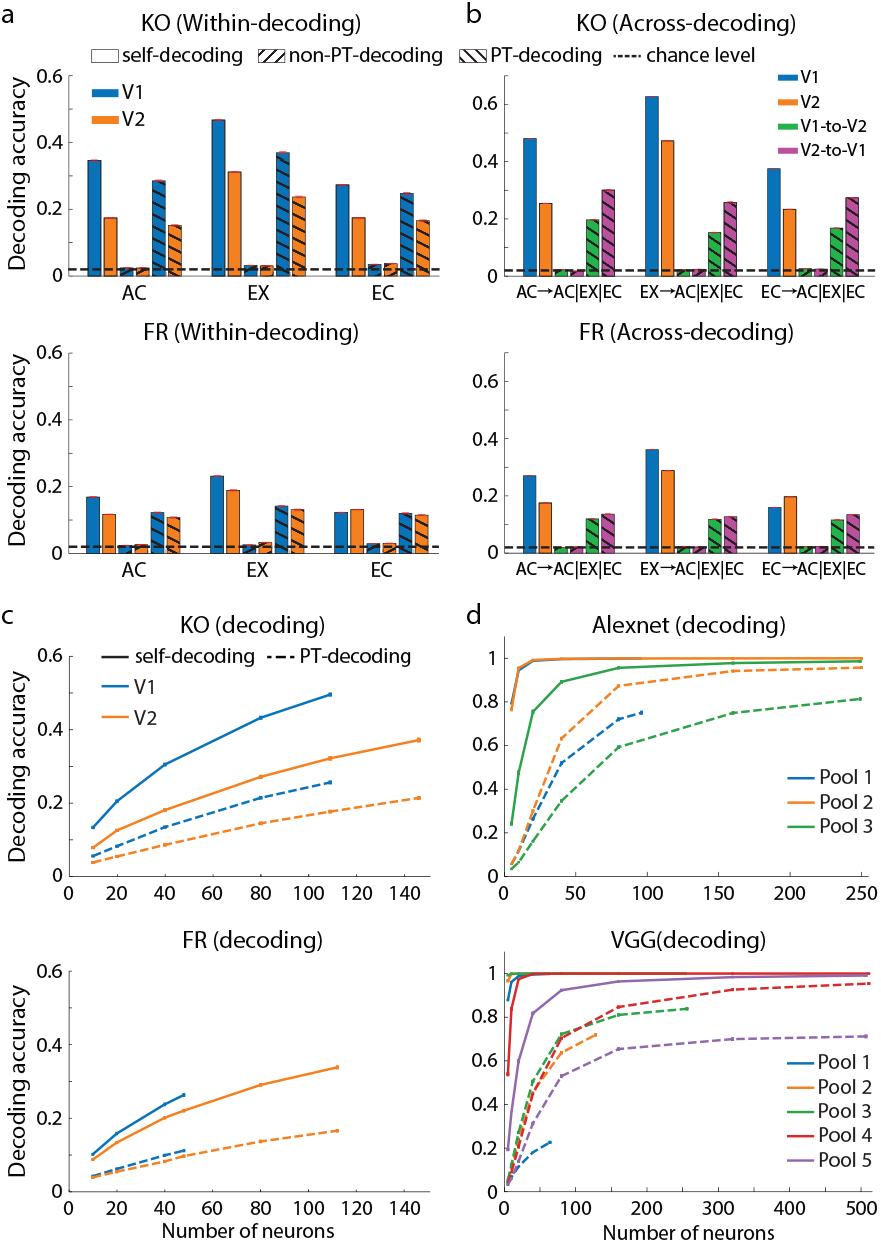
Cue-transfer population decoding within and across regions and effect of population size. (a) Within-region transfer decoding. Decoding accuracies were estimated between two different subpopulations of neurons from the same region responding to the same rendering (monkey KO, top; FR, bottom). Blue and orange bars indicate V1 and V2 data, respectively. The x-axis indicates the stimulus rendering. For each rendering, bars show self-decoding (plain bars, an upper bound), non-PT-decoding (forward-slash hatching, transfer decoding without PT), and PT-decoding (backslash hatching, transfer decoding with PT). The dashed line indicates the 2% chance level. Error bars are SEM. (b) Across-region transfer decoding. Decoding accuracies were estimated for both V1-to-V2 (train on V1, test on V2) and V2-to-V1 (train on V2, test on V1) transfers in monkey KO (top) and FR (bottom). The x-axis notation, in the format Ttrain → Test1|Test2|Test3, indicates that the plotted bar is the average decoding accuracy across the three specified test renderings using the same training rendering. Bar colors denote V1 self-decoding (blue), V2 self-decoding (orange), V1-to-V2 decoding (green), and V2-to-V1 decoding (magenta). Bar styles (plain, hatched, etc.) are as in (a). (c) Effect of population size on neural decoding. Accuracy is plotted as a function of the number of neurons for V1 (blue) and V2 (orange). Solid lines represent self-decoding and dashed lines represent PT cue-transfer decoding. Horizontal bars on each data point show the SEM. (d) Effect of population size on model decoding. Accuracy is plotted as a function of population size for the pooling layers of AlexNet (top) and VGG (bottom). Panel details are as in (c).

**Figure 5:**
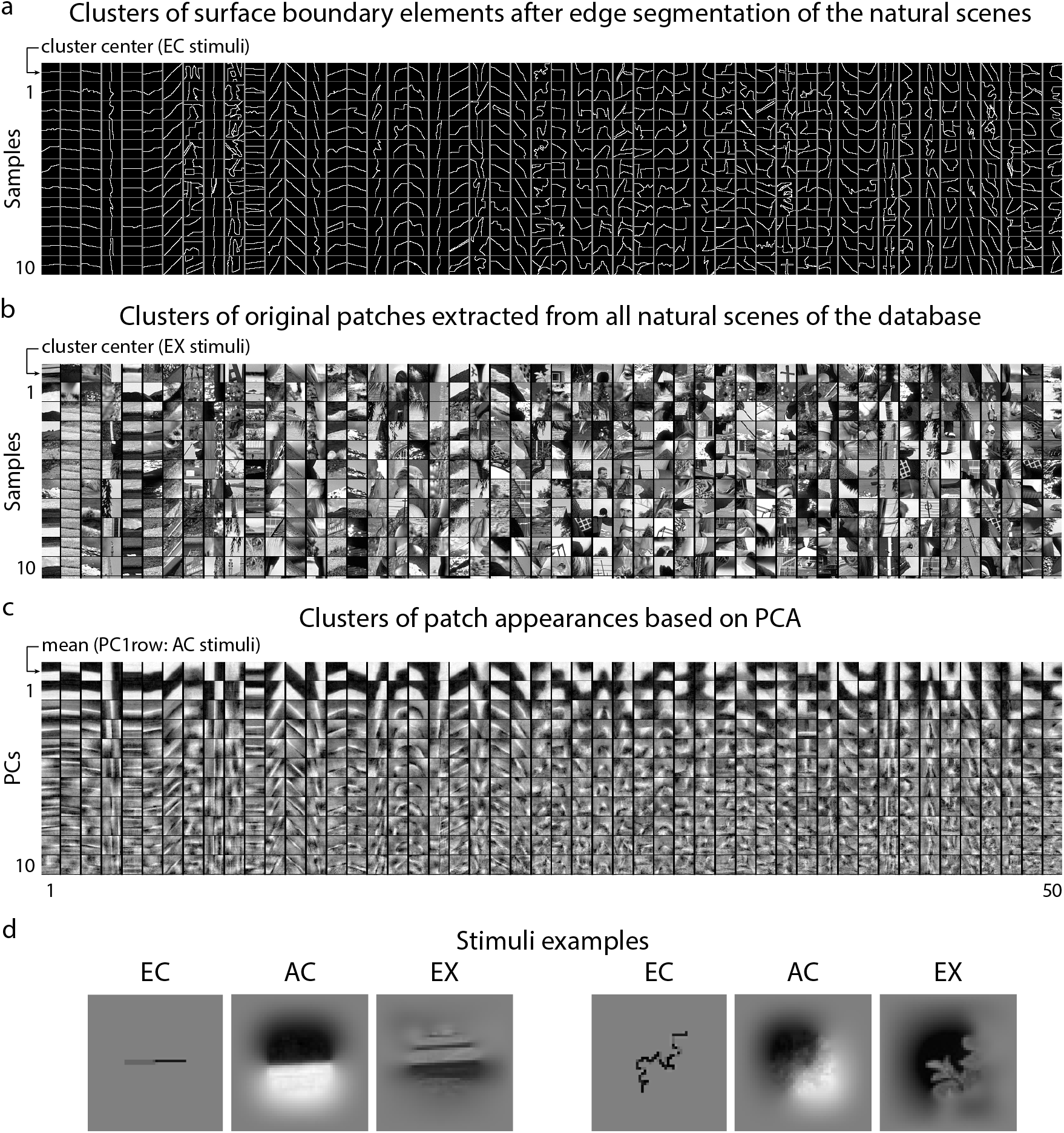
Dataset of surface boundary stimuli. (a) Clusters of surface boundary elements after edge segmentation of the natural scenes of the Berkeley segmentation database. For each of the 50 clusters, 10 example samples of surface-boundary elements are shown in the numbered rows (1-10). The top ‘cluster center’ row displays the segmented surface-boundary element at the center of each cluster; these 50 elements form the ‘Edge Contours’ (EC) dataset. (b) Clusters of the original patches extracted for the natural scenes corresponding to the clusters in (a). Following the same layout, the top ‘cluster center’ row displays the patch at the center of each cluster; these 50 patches form the ‘Example’ (EX) dataset. (c) Clusters of patch appearances derived from PCA. For each cluster, the top row displays the mean patch for reference. The subsequent 10 rows (labeled 1-10) display the first 10 eigen-images (PCs) for each cluster. The first of these numbered rows (PC1) forms the ‘Appearance Contours’ (AC) dataset and is visually similar to the mean patch in the top row. (d) Two stimulus examples, each shown in its three renderings: Edge Contour (EC), Appearance Contour (AC), and Example (EX).

To do so, we measured a ‘tuning correlation’ by computing the Pearson correlation between each pair of tuning curves (i.e., EC-AC, EC-EX, and AC-EX). Fig.2.c shows the tuning correlations for the example V1 neuron, which had a significant correlation for all three rendering pairs (*p <* 0.01), indicating some degree of cue-invariance across all renderings. We then measured the tuning correlations of all neurons in V1 (Fig.2.d) and V2 (Fig.8.a) and quantified the proportion of neurons that showed significant tuning correlations for one-, two-, and three-pair rendering sets (see the caption of Fig.2 for the full list of correlation sets). In monkey KO, the proportion of V1 neurons showing significant tuning correlation (i.e., ‘cue-invariance’) was, on average, 34%, 25%, and 22% for one-, two-, and three-pair rendering sets, respectively. These proportions were all significantly above chance, as the lower bound of the Wilson score interval exceeded the chance level for all correlation sets (Table 1, highlighted row ‘V1/W.(lower)’). In monkey FR, the results were similar, although the proportions of cue-invariant neurons were lower. This may be due to differences in monkeys’ visual experiences and training. The proportion of V1 neurons with significant correlations was, on average, 22%, 10%, and 6% for one-, two-, and three-pair rendering sets, respectively, and these proportions were also significantly above chance (see Tables 1 and 2 for a detailed description of the statistical results for monkey KO and FR). The proportion of V2 neurons showing cue-invariance was lower than in V1 for both animals. For monkey KO, the corresponding proportions were, on average, 14%, 6%, and 5%, and were significantly above chance (Wilson score interval; see Table 1, highlighted row ‘V2/W.lower’). For monkey FR, the corresponding proportions were on average 13%, 5%, and 4%, and these proportions were also significantly above chance (Wilson score interval; see Table 2, highlighted row ‘V2/W.lower’). Notably, for both monkeys and brain areas, the significant tuning correlations corresponded to the highest correlation levels (Fig.7). The proportion of neurons showing cue-invariance was significantly higher in V1 than in V2 for all correlation sets for monkey KO (CPP, *p <* 0.01; highlighted row ‘V1vsV2/CPP(p)’ in Table 1). In monkey FR, this difference was only significant for all one-pair correlation sets and for the specific two-pair set ‘2a’ (EC-AC and EC-EX) (CPP, *p <* 0.01; highlighted row ‘V1vsV2/CPP(p)’ in Table 2).

**Table 1:** Cue-invariant statistics of V1 and V2 neurons for monkey KO. ‘%’ and ‘N’ indicate the proportion and number of neurons with significant tuning correlation for each correlation set. ‘SSE’ indicates the standard error of the sample proportion and is used for the error bars in the main figures. ‘W. lower’ and ‘W. upper’ indicate the lower and upper bounds of the Wilson score interval, respectively. The chance levels for one-(e.g., ‘ec-ac’), two-(e.g., ‘ec-ac and ec-ex’), and three-pair (‘ec-ac-ex’) correlation sets are *p* = 0.05, *p* = 2.5 × 10^−3^, and *p* = 1.25 × 10^−4^, respectively. Cells highlighted in yellow indicate that the lower bound of the Wilson score interval was higher than the chance level. Cells highlighted in green indicate that the p-value of the Comparison of Population Proportions (CPP) was significant (*p <* 0.01).

| monkey KO |  | Correlation sets |  |  |  |  |  |  |  |
| --- | --- | --- | --- | --- | --- | --- | --- | --- | --- |
|  |  | ec-ac | ec-ex | ac-ex | ec-ac-ec-ex | ec-ac-ac-ex | ec-ex-ac-ex | ec-ac-ex | all |
| V1 | % | 0.33945 | 0.42202 | 0.27523 | 0.26606 | 0.22936 | 0.25688 | 0.22018 | 1 |
|  | Mean | 0.34556 |  |  | 0.25076 |  |  | 0.22018 | 109 |
|  | N | 37 | 46 | 30 | 29 | 25 | 28 | 24 |  |
|  | SSE | 0.04535 | 0.04730 | 0.04277 | 0.04232 | 0.04026 | 0.04184 | 0.03968 |  |
|  | W.(lower) | 0.25737 | 0.33350 | 0.20011 | 0.19209 | 0.16045 | 0.18411 | 0.15266 |  |
|  | W.(upper) | 0.43245 | 0.51583 | 0.36564 | 0.35594 | 0.31668 | 0.34619 | 0.30675 |  |
| V2 | % | 0.16438 | 0.10959 | 0.14384 | 0.04794 | 0.06849 | 0.06164 | 0.04794 | 1 |
|  | Mean | 0.13927 |  |  | 0.05936 |  |  | 0.04794 | 146 |
|  | N | 24 | 16 | 21 | 7 | 10 | 9 | 7 |  |
|  | SSE | 0.03067 | 0.02585 | 0.02904 | 0.01768 | 0.02090 | 0.01990 | 0.01768 |  |
|  | W.(lower) | 0.11302 | 0.06858 | 0.09604 | 0.02341 | 0.03762 | 0.0327 | 0.02341 |  |
|  | W.(upper) | 0.23295 | 0.17060 | 0.20989 | 0.09565 | 0.12148 | 0.11299 | 0.09565 |  |
| V1 vs. | CPP(z) | 4.2321 | 7.5528 | 3.1764 | 5.2727 | 3.8888 | 4.7197 | 4.1637 |  |
| V2 | CPP(p) | 1.16E-05 | 2.13E-14 | 7.45E-04 | 6.72E-08 | 5.04E-05 | 1.18E-06 | 1.57E-05 |  |

**Table 2:** Cue-invariant statistics of V1 and V2 neurons for monkey FR. ‘%’ and ‘N’ indicate the proportion and number of neurons with significant tuning correlation for each correlation set. ‘SSE’ indicates the standard error of the sample proportion and is used to plot the error bars in the main figures. ‘W. lower’ and ‘W. upper’ indicate the lower and upper bounds of the Wilson score interval, respectively. The chance levels for one-(e.g., ‘ec-ac’), two-(e.g., ‘ec-ac and ec-ex’), and three-pair (‘ec-ac-ex’) correlation sets are *p* = 0.05, *p* = 2.5× 10^−3^, and *p* = 1.25 ×10^−4^, respectively. Cells highlighted in yellow indicate that the lower bound of the Wilson score interval was higher than the chance level. Cells highlighted in green indicate that the p-value of CPP was significant (*p <* 0.01).

| monkey FR |  | Correlation sets |  |  |  |  |  |  |  |
| --- | --- | --- | --- | --- | --- | --- | --- | --- | --- |
|  |  | ec-ac | ec-ex | ac-ex | ec-ac-ec-ex | ec-ac-ac-ex | ec-ex-ac-ex | ec-ac-ex | all |
| V1 | % | 0.27083 | 0.20833 | 0.1875 | 0.16667 | 0.08333 | 0.0625 | 0.0625 | 1 |
|  | Mean | 0.22222 |  |  | 0.10416 |  |  | 0.0625 | 48 |
|  | N | 13 | 10 | 9 | 8 | 4 | 3 | 3 |  |
|  | SSE | 0.06414 | 0.05861 | 0.05633 | 0.05379 | 0.03989 | 0.03493 | 0.03493 |  |
|  | W.(lower) | 0.16565 | 0.11730 | 0.10191 | 0.08695 | 0.03288 | 0.02148 | 0.02148 |  |
|  | W.(upper) | 0.40997 | 0.34259 | 0.31940 | 0.29578 | 0.19553 | 0.16835 | 0.16835 |  |
| V2 | % | 0.16964 | 0.11607 | 0.09821 | 0.0625 | 0.0625 | 0.03571 | 0.03571 | 1 |
|  | Mean | 0.12797 |  |  | 0.05357 |  |  | 0.03571 | 112 |
|  | N | 19 | 13 | 11 | 7 | 7 | 4 | 4 |  |
|  | SSE | 0.03546 | 0.03026 | 0.02812 | 0.02287 | 0.02287 | 0.01753 | 0.01753 |  |
|  | W.(lower) | 0.11137 | 0.06909 | 0.05572 | 0.03060 | 0.03060 | 0.01397 | 0.01397 |  |
|  | W.(upper) | 0.24981 | 0.18850 | 0.16734 | 0.12341 | 0.12341 | 0.08824 | 0.08824 |  |
| V1 vs. V2 | CPP(z) | 2.8677 | 2.6147 | 2.5303 | 2.952 | 0.59041 | 0.7591 | 0.7591 |  |
|  | CPP(p) | 0.0020 | 0.00446 | 0.00569 | 0.00157 | 0.27746 | 0.2239 | 0.2239 |  |

To gain insights into the cortical processing underlying cue-invariance, we first grouped V1 and V2 neurons by their cortical layer (Vanni et al., 2020). We estimated the laminar positions of each neuron using a combination of Current-Source Density analysis and the onset latency of the visual activity across the contacts of the laminar probe (see Methods, Fig.6). Neurons were grouped into 3 main ensembles, namely: ‘Superficial’, ‘Layer 4’, and ‘Deep’ layers (Fig.6.a). We then recomputed the proportion of neurons with significant tuning correlations within each layer of V1 and V2 for both monkeys (Fig.2.e). The main observation, consistent across both animals, was a higher proportion of V1 neurons within Layer 4 showing cue-invariance compared to the other layers. In monkey KO, the average proportion of V1 neurons in Layer 4 was higher for all correlation sets, but significant only for the EC-EX (1b) pair (on average, 67% of V1 neurons; CPP: Layer 4 vs. Superficial layers, *p* = 3.5 × 10^−4^; Layer 4 vs. Deep Layers, *p* = 7.8 × 10^−4^). For monkey FR, the average proportion of V1 neurons in Layer 4 was also higher for several correlation sets (EC-AC (1a), EC-EX (1b), and EC-AC and EC-EX (2a)). Like in monkey KO, it was significant for the EC-EX (1b) pair (on average, 35% of V1 neurons; CPP: Layer 4 vs. Superficial Layers, *p* = 2.5 × 10^−3^; Layer 4 vs. Deep Layers, *p* = 9.2 × 10^−4^). Other layer-specific patterns were not consistent across animals. For instance, in monkey FR, Superficial Layers lacked a significant proportion of cue-invariant V1 neurons, while Deep Layers showed a proportion that was equal to or higher than that of Layer 4. The same patterns were not observed in monkey KO. In V2, we found no consistent pattern of cue-invariance across the cortical layers for either monkey (Fig.8.a). In a separate analysis, we investigated whether cue-invariance depended on a specific temporal window, by analyzing an early neural activity time window (40-110 ms) and a late activity related to putative feedback from downstream regions (110-180 ms) (see Methods, Fig.6.d). This revealed that in V1, the proportion of cue-invariant neurons did not depend on the temporal window of analysis (Fig.8.b,c). In V2, however, the results were inconsistent across monkeys.

**Figure 6:**
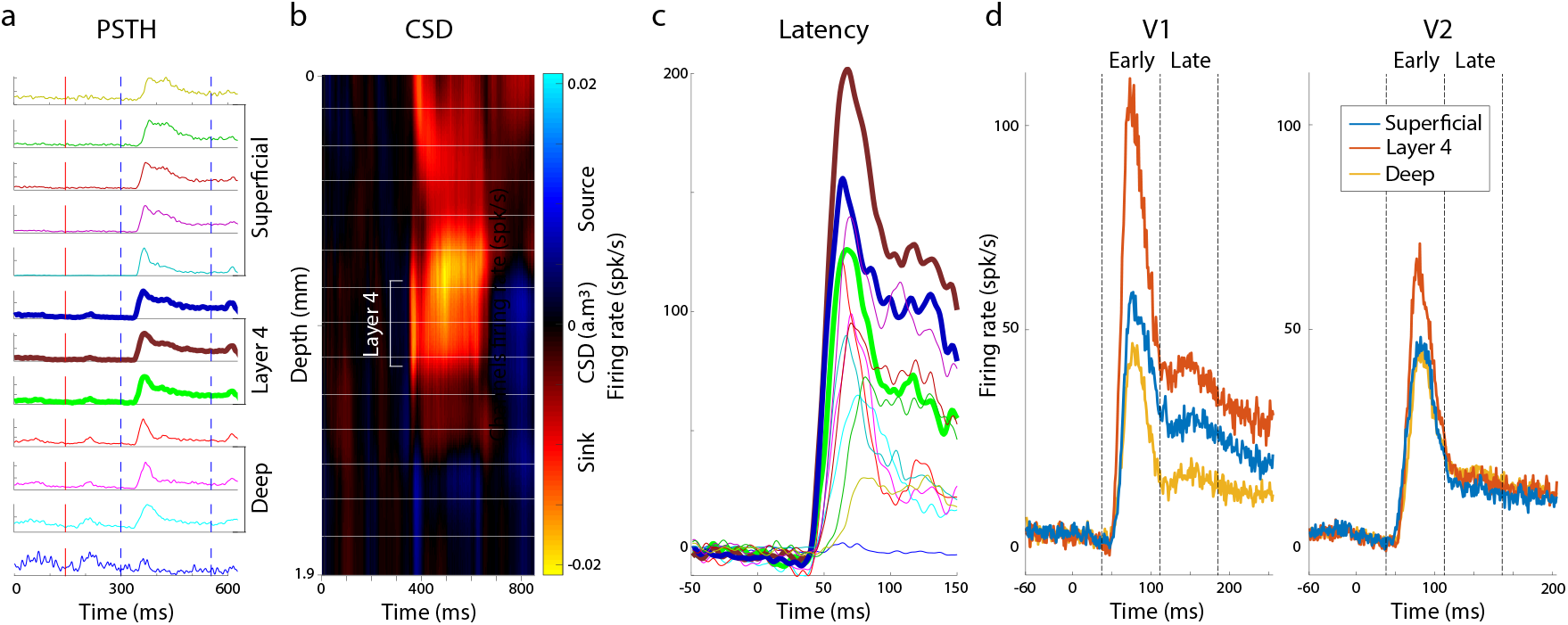
Neurophysiological recording. (a) Example of a recording session with a linear multi-contact probe (only 12 contacts shown). Each row displays the PSTHs recorded on each channel after offline spike sorting. Channels are grouped into layers (Deep, Layer 4, and Superficial) based on the current source density (CSD) and onset latency information (see Methods). (b) CSD associated with the example recording session in (a). The ‘Layer 4’ label indicates the location of the input source signal. (c) All PSTHs from the example session in (a), aligned on stimulus onset and used for onset latency estimation. (d) Trial-averaged firing rate across all sessions and neurons in V1 and V2, grouped by their estimated layers. Vertical black dashed lines indicate the limits of the temporal windows used to define early and late responses.

**Figure 7:**
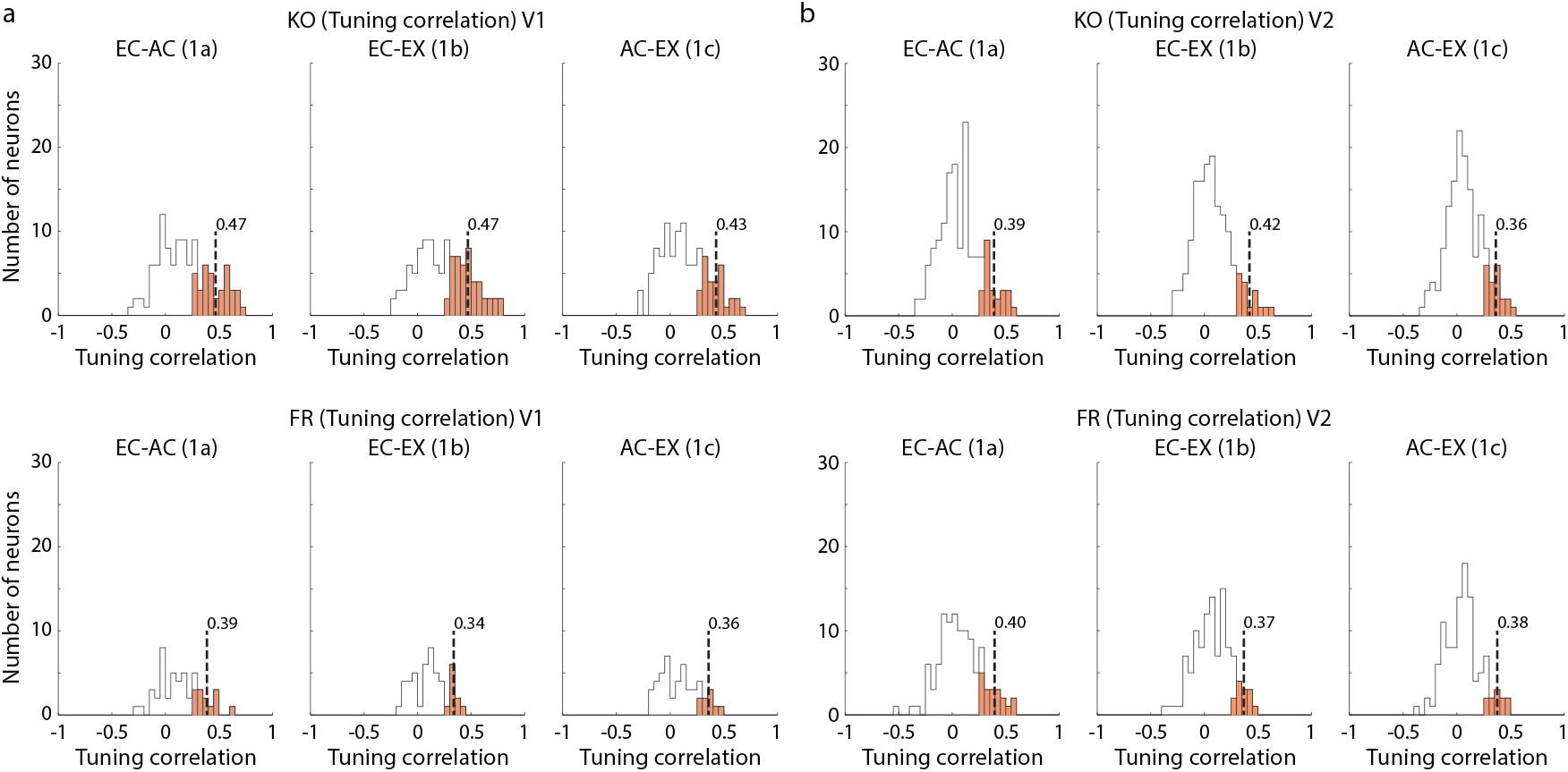
Distributions of tuning correlations of individual V1 and V2 neurons for the different correlation sets. (a) Distributions of tuning correlations for individual V1 neurons in monkey KO (top) and FR (bottom) across three rendering pairs (EC-AC (1a); EC-EX (1b); AC-EX (1c)). The black outline indicates the full distribution of correlations. The filled orange bars show the distribution of only the significant positive correlations. The vertical dashed line indicates the mean of this significant distribution, which corresponds to the set of neurons used for the results presented in Fig.2.d. (b) Distributions of tuning correlations for individual V2 neurons in monkey KO (top) and FR (bottom). Panel details are as in (a).

**Figure 8:**
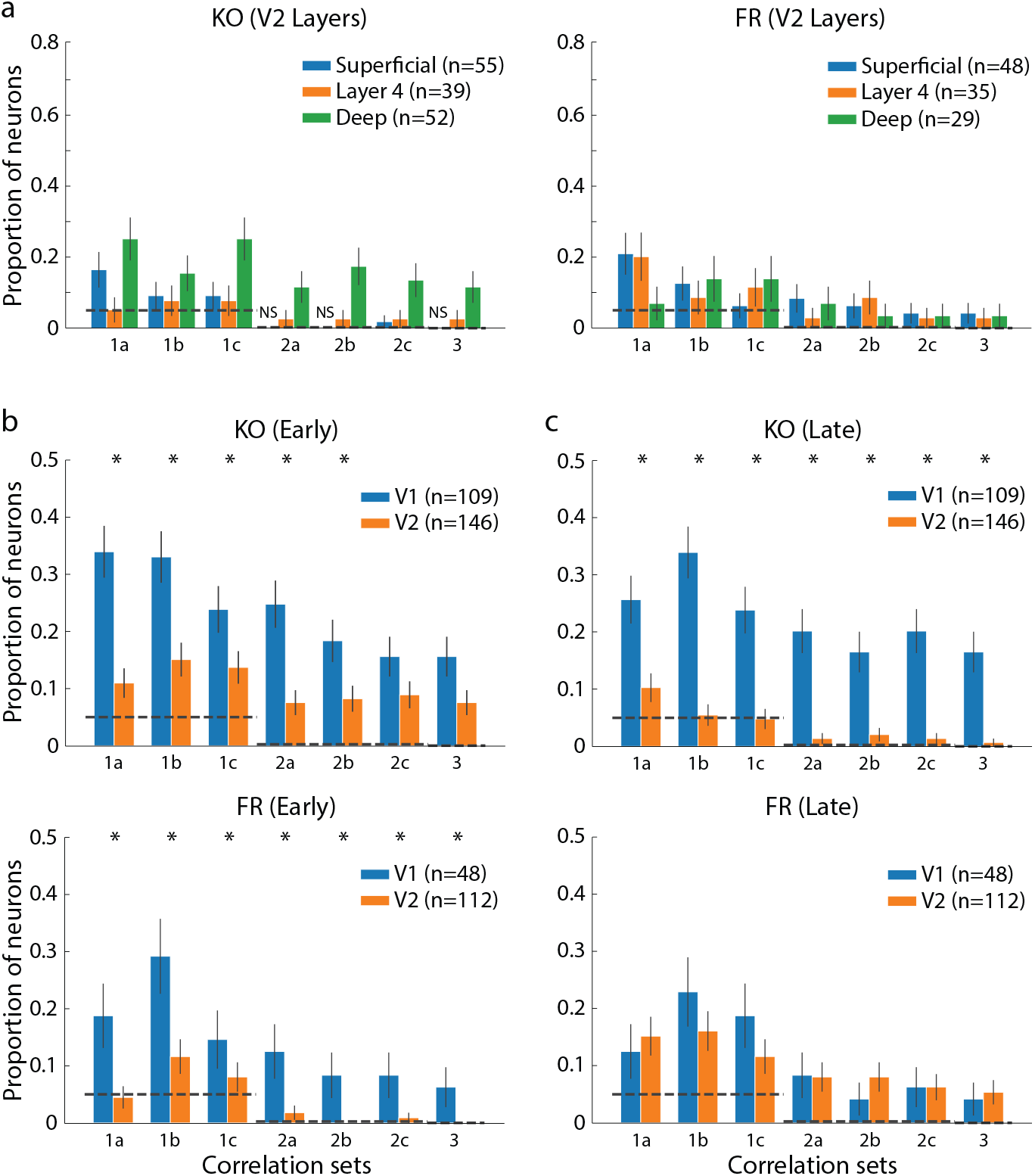
Layer and temporal window analysis of tuning correlations. (a) Proportion of cue-invariant V2 neurons grouped by cortical layer: Superficial (blue), Layer 4 (orange), and Deep (green), for monkey KO (left) and FR (right). The dashed line indicates the chance level. (b, c) Proportion of cue-invariant V1 (blue) and V2 (orange) neurons for early (b) and late (c) response windows, shown for monkey KO (top) and FR (bottom). Dashed lines indicate the chance level for one-(*p* = 0.05), two-(*p* = 2.5 × 10^−3^), and three-pair (*p* = 1.25 × 10^−4^) correlation sets. Asterisks (*) indicate a significant difference between V1 and V2. For all panels, bars indicate the proportion of neurons with significant tuning correlations for each correlation set.

Overall, a significant proportion of individual neurons in V1 and V2 showed cue-invariance across renderings. This effect was more pronounced in V1, where neurons were proportionally more cue-invariant than in V2; in particular, a significant fraction of V1 neurons showed cue-invariance across multiple rendering pairs. These findings suggest that early visual areas like V1 may represent basic visual elements, such as oriented edges, in a more cue-invariant manner (see Discussion). However, the cue-invariance measured through tuning correlations of individual neurons was limited. Therefore, we next analyzed the cue-invariance properties at the population level.

### Cue-Invariant Properties of the Geometric Structure of V1 and V2 Population Codes

Neural population analyses can complement single-neuron approaches and improve our understanding of information processing in the brain (Kriegeskorte and Kievit, 2013; Kriegeskorte et al., 2008; Yan et al., 2014). Here, we analyzed the cue-invariance properties of V1 and V2 population responses to examine whether they could reveal stronger invariance than was observed at the single-neuron level.

First, we applied representational similarity analysis (RSA) to population response patterns for each pair of renderings (see Methods). RSA is a general, model-agnostic method for characterizing the information content of population activity (Kriegeskorte et al., 2008). It makes no assumptions about readout/decoding mechanisms or about the structure (geometry) of the population code (Kriegeskorte and Kievit, 2013). While this revealed similarity for some stimulus pairs, the results were inconsistent across animals (Fig.9). To access information about the population structure not captured by RSA, we therefore turned to an alternative, decoding-based analysis.

**Figure 9:**
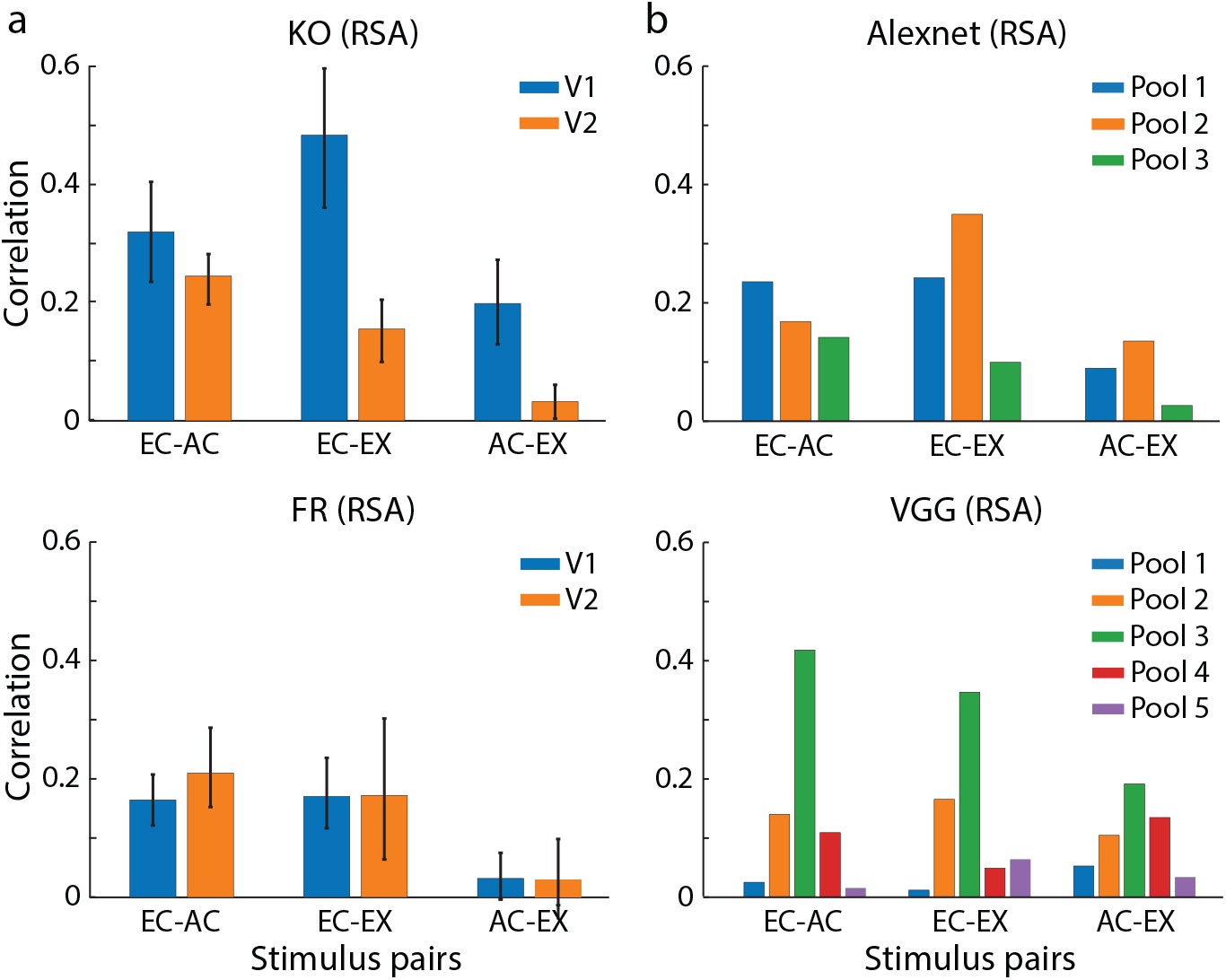
Representation similarity analysis (RSA). (a) RSA results for V1 (blue) and V2 (orange) in monkey KO (top) and FR (bottom). The x-axis indicates the rendering pairs used for the analysis. Error bars indicate the 95% confidence interval. (b) RSA results for the pooling layers of AlexNet (top) and VGG (bottom). The format is the same as in panel (a), but the bar colors correspond to the different pooling layers as indicated in the legend.

To assess the cue-invariant characteristics of the population codes, we developed a ‘cue-transfer’ decoding method (see Methods). In this context, we use ‘renderings’ and ‘cues’ interchangeably. In brief, a linear support vector machine (SVM) was trained on the z-scored neural responses (using a 300-ms window from 30 ms to 330 ms after stimulus onset) to visual stimuli rendered in one cue and tested on its accuracy in decoding the same stimuli rendered in a different cue (see the ‘transfer decoding’ diagram in Fig.10.b). To control for the effect of population size, the V2 sample size was matched to that of V1 for each monkey in all tests. The decoding procedure was performed reciprocally for each cue pair (e.g., for the AC-EC pair, training on AC and testing on EC, and vice versa), and the final decoding accuracy was the average of these two estimations (Fig.3.b; results labeled ‘non-PT-decoding’ denote transfer decoding without the Procrustes Transformation, which is introduced below).

Principal component analysis (PCA) of the population responses for each rendering reveals a consistent geometric arrangement of the stimulus representations (see 2D PCA projection in Fig.3.a). This consistency suggested that a shared representational geometry may underlie the different cues and that geometric transformations could be used to align the cue-specific population codes. To test this hypothesis, we used Procrustes Transformation (PT) to find the optimal transformation that aligns the population codes from two different cues (i.e., the optimal rotation, translation, and scaling). In this method, the trial-averaged population response to each stimulus served as ‘landmarks’ to align the population codes of two cues using PT. Specifically, the goal was to align the set of landmarks from one rendering (the ‘comparison shape’) to another (the ‘target shape’; see Methods and Fig.10). Fig.3.a illustrates this process, showing how the population codes from the three renderings align more closely after the transformations are applied (second row of Fig.3.a). Finally, cue-transfer decoding was performed on the Procrustes-aligned population codes (Fig.3.b; results labeled ‘PT-decoding’).

**Figure 10:**
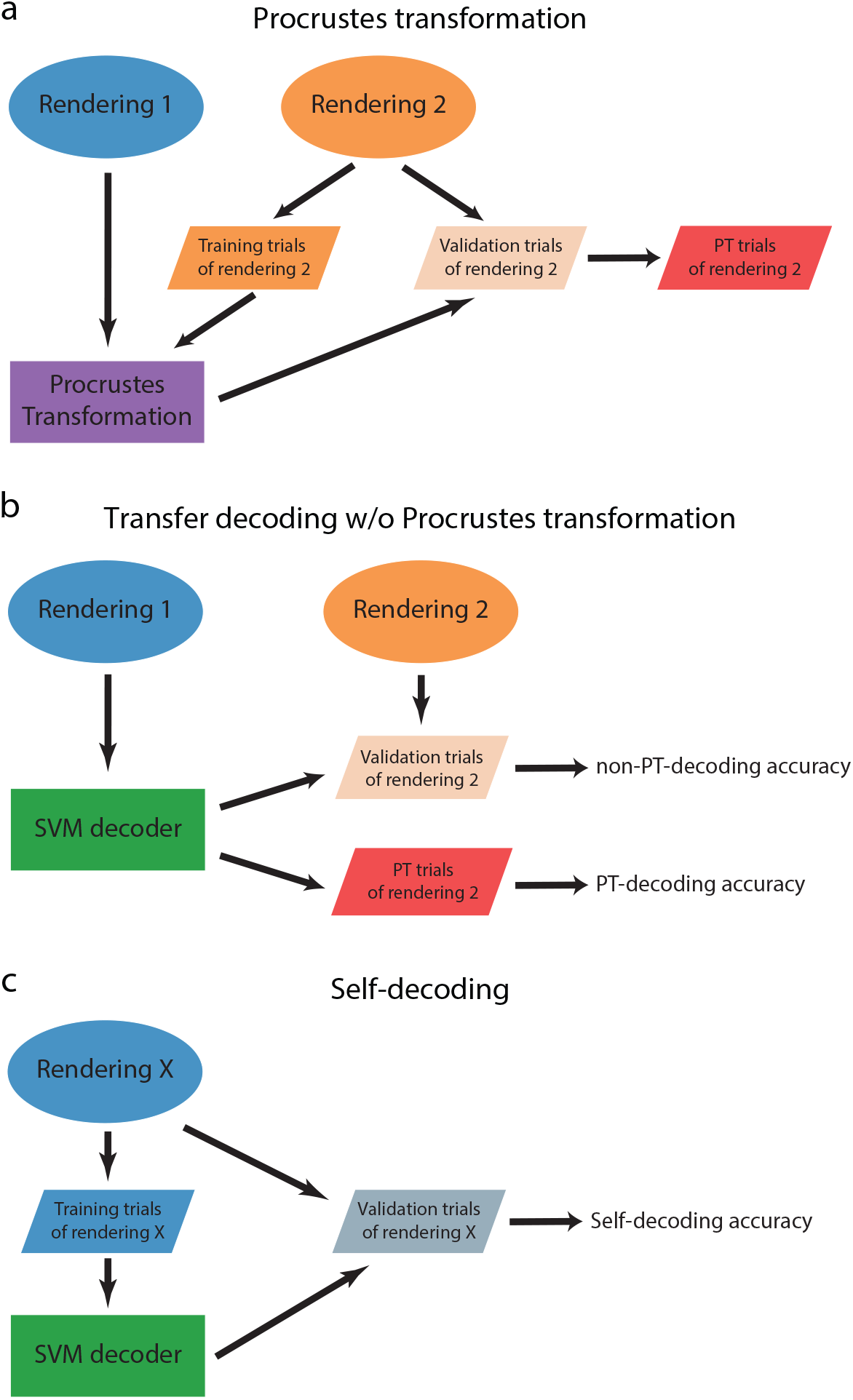
Transfer-decoding diagram. (a-c) Each diagram illustrates the scheme used to estimate decoding accuracy. (a) Procrustes Transformation. The transformation is calculated using the full population response to Rendering 1 (target) and the training trials from Rendering 2 (comparison). This transformation is then applied to the held-out validation trials of Rendering 2 to generate the transformed ‘PT trials of rendering 2.’ (b) Transfer Decoding. An SVM decoder is trained on the responses to Rendering 1. It is then tested on both the original validation trials from Rendering 2 (to get non-PT accuracy) and the transformed ‘PT trials’ (to get PT accuracy). (c) Self-Decoding. An SVM decoder is trained on the training trials of a given rendering (Rendering X) and tested on the validation trials of the same rendering to establish an upper-bound performance.

To benchmark this cue-transfer decoding performance, we established upper and lower bounds (see Methods). The upper bound, or ‘self-decoding’ accuracy, was estimated by training and testing the decoder on population responses to the same rendering (‘self-decoding’ diagram, Fig.10.c). The theoretical lower bound was the 2% chance level for decoding 50 stimuli (indicated by the black dashed lines on Fig.3.b-d). We empirically confirmed this chance level by shuffling the stimulus labels when performing PT. This shuffling removes the one-to-one stimulus correspondence between the ‘target’ and ‘comparison’ landmark matrices, each consisting of trial-averaged population responses to one of the two different renderings (see Methods for a detailed description, and ‘shuffle of stimulus labels’ in Fig.11.a). This procedure effectively reduced the decoding accuracies to the theoretical chance level (Fig.11.b).

**Figure 11:**
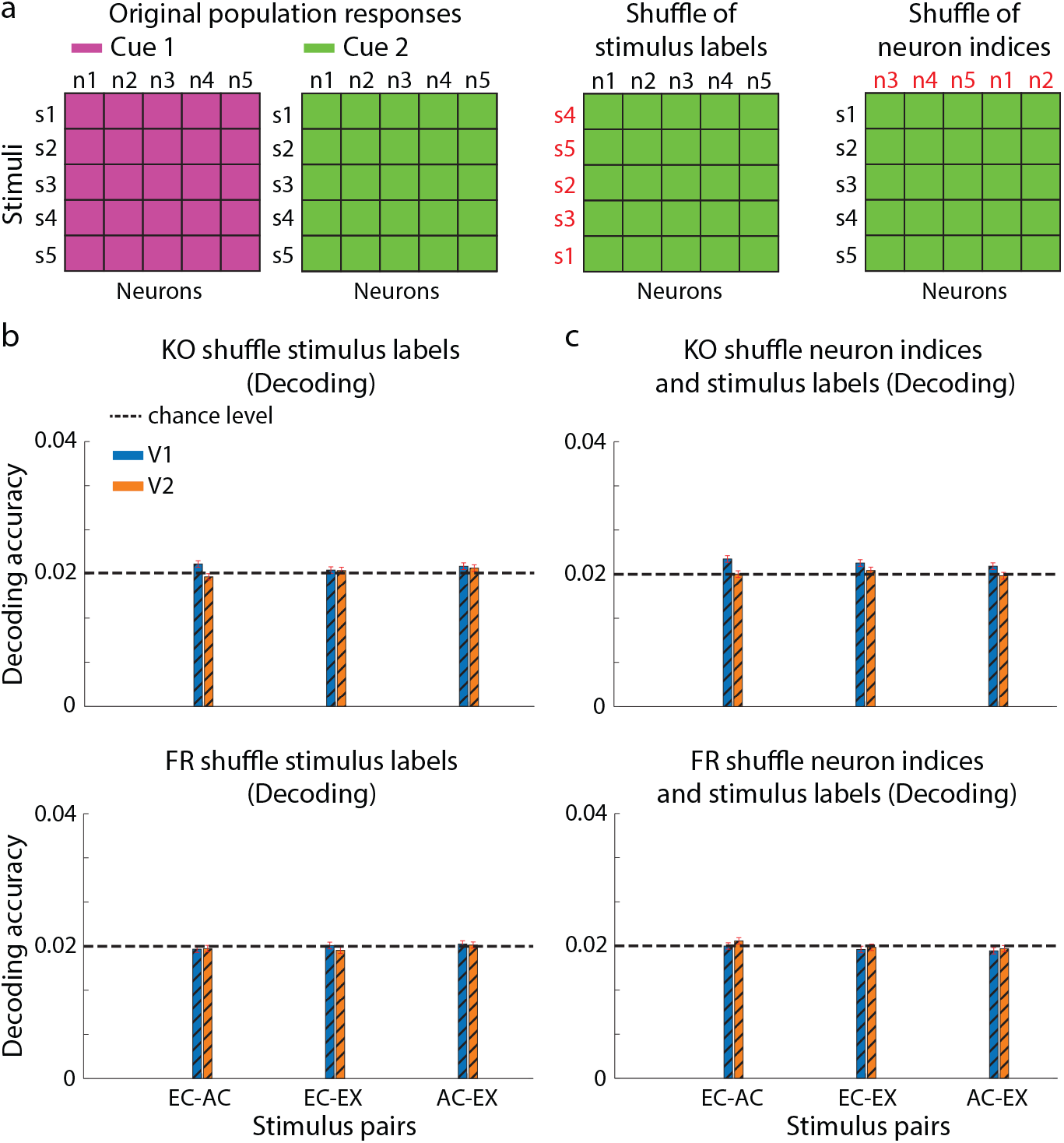
Population shuffle procedures and estimations. (a) Diagram of the shuffle procedures. Left: The original population responses for two cues. Middle: The ‘shuffle of stimulus labels’ procedure, where the stimulus labels (rows) for Cue 2 are randomized. Right: The ‘shuffle of neuron indices’ procedure, where the neuron identities (columns) for Cue 2 are randomized. (b) Population decoding accuracy after applying the stimulus label shuffle in PT. Results are shown for monkey KO (top) and FR (bottom). Blue and orange bars indicate V1 and V2 data, respectively. The horizontal dashed line indicates the 2% chance level. Red error bars indicate SEM. (c) Decoding performance after applying both a neuron index shuffle to the original data matrices and a stimulus label shuffle in PT. Panel details are as in (b).

Fig.3.b shows the cue-transfer decoding performance for V1 and V2 populations in monkeys KO and FR, both before and after applying Procrustes Transformation. For both V1 and V2, decoding accuracy in the non-PT and PT conditions was well above the chance level, indicating that population codes in each area had some degree of cue-invariance. Applying the Procrustes Transformation markedly improved cue-transfer decoding, yielding accuracies two-to fourfold higher than the non-PT decoding accuracy in both V1 and V2 and for both animals (for monkey KO, PT accuracy was, on average, 323% of the non-PT level [95% CI: 317%, 330%] in V1 and 425% [95% CI: 416%, 434%] in V2; for monkey FR, the corresponding values were 314% [95% CI: 308%, 321%] in V1 and 269% [95% CI: 263%, 275%] in V2). While PT decoding accuracy was still lower than the ‘self-decoding’ upper bound, the transformation recovered approximately half of the decodable information across renderings (for monkey KO, PT accuracy was, on average, 56% of the self-decoding level [95% CI: 55%, 57%] in V1 and 64% [95% CI: 63%, 65%] in V2; for monkey FR, the corresponding values were 49% [95% CI: 48%, 50%] in V1 and 47% [95% CI: 46%, 48%] in V2). This indicates that the geometric structure of the neural data, when transformed appropriately, can be used to recover cue-invariant properties of the visual inputs across different renderings.

PT cue-transfer decoding accuracy was consistently higher in V1 than in V2 for both monkeys (for monkey KO, V1 PT accuracy was, on average, 7.86 percentage points higher than V2 [95% CI: 7.63, 8.09]; for monkey FR, the corresponding value was 1.65 percentage points [95% CI: 1.48, 1.83]). Similar to our single-neuron tuning findings, this result indicates that V1 populations show more cue-invariance than V2 populations, meaning the geometry of the population code is better preserved across cues in V1. This advantage may be explained by the V1 population code being more discriminative of the surface boundary elements within each rendering. This hypothesis is also supported by the self-decoding accuracies, which were also consistently higher in V1 than in V2 for both monkeys (for monkey KO, V1 was higher by a mean of 17.68 percentage points [95% CI: 17.55, 17.82]; for monkey FR, the corresponding value was 4.25 percentage points [95% CI: 4.06, 4.44]).

While non-PT cue-transfer decoding may rely on the tuning correlations of individual neurons, PT decoding leverages the geometric structure of the population code and may not depend on the tuning correlation of individual neurons. To evaluate the contribution of the tuning correlation to the cue-transfer decoding, we performed a neuron-shuffling experiment. Specifically, we randomized the neuron indices between the two population response matrices before applying PT, a procedure that preserves the overall population geometry while breaking the neuron-by-neuron tuning correspondence across cues (see Methods and Fig.11.a). This shuffle reduced non-PT cue-transfer decoding to chance levels but left PT cue-transfer decoding performance largely unaffected (Fig.3.c). These results demonstrate that eliminating single-neuron tuning correlations across cues does not adversely affect PT cue-transfer decoding, suggesting that cue-invariance is primarily encoded by the geometric structure of the population code, which can supplement or even supersede the role of single-neuron tuning.

### Invariant Geometric Structures within and across Regions

In the previous section, we established that the geometric structure of V1 and V2 populations encodes cue-invariant properties. Here, we investigate whether these properties are also preserved in distinct neuronal subpopulations, both within and across these two areas.

To further characterize the nature of this geometric invariance, we next tested whether the structure was stable not only across cues but also across different subsets of neurons responding to the same cue. In this ‘within-decoding’ analysis (see Methods), we randomly split the recorded population from a single area into two non-overlapping subpopulations, and tested if a decoder trained on one could read out responses from the other to the same stimulus rendering (Fig.4.a). As expected, non-PT transfer decoding remained at chance level, since there was no direct neuron-to-neuron tuning correspondence between the two arbitrary subpopulations. However, after applying PT to align the two subpopulation codes, transfer decoding accuracies approached the self-decoding upper bounds (for monkey KO, PT accuracy was, on average, 86.29% of the self-decoding level [95% CI: 85.20%, 87.51%] in V1 and 89.89% [95% CI: 88.35%, 91.45%] in V2; for monkey FR, the corresponding values were 80.60% [95% CI: 78.96%, 82.32%] in V1 and 87.42% [95% CI: 85.56%, 89.29%] in V2). Crucially, this result shows that the population geometry is stable across different neuronal subsets, demonstrating a ‘neuron-invariant’ coding property. This supports the broader hypothesis that the brain can achieve robust representations that are invariant to both the visual cue and the specific neurons being read out.

To explore whether the invariant geometric structure is shared between V1 and V2, we applied our cue-transfer decoding method on populations across the two areas (see ‘across-decoding’ in Methods). We performed two reciprocal analyses: ‘V1-to-V2’ decoding, where a decoder trained on V1 population responses was tested on V2, and ‘V2-to-V1’ decoding, which was the reverse. We applied the transfer decoding procedure to stimuli of the same rendering and of different renderings (for a detailed description of all decoding scenarios, see Fig.12). Fig.4.b shows the average accuracies of V1-to-V2 and V2-to-V1 decoding. Both V1-to-V2 and V2-to-V1 non-PT transfer decoding accuracies were around chance levels, due to the lack of a direct neuron-by-neuron tuning correspondence between the V1 and V2 populations. On the contrary, cross-area PT decoding accuracies were significantly above chance levels. The substantial improvement provided by PT was comparable to that observed in the within-region analysis (Fig.4.a), demonstrating that the transformation is effective at aligning population codes both within a region and across different brain areas. These findings suggest that a shared, invariant geometric structure is a plausible mechanism for encoding cue-invariant properties in V1 and V2.

**Figure 12:**
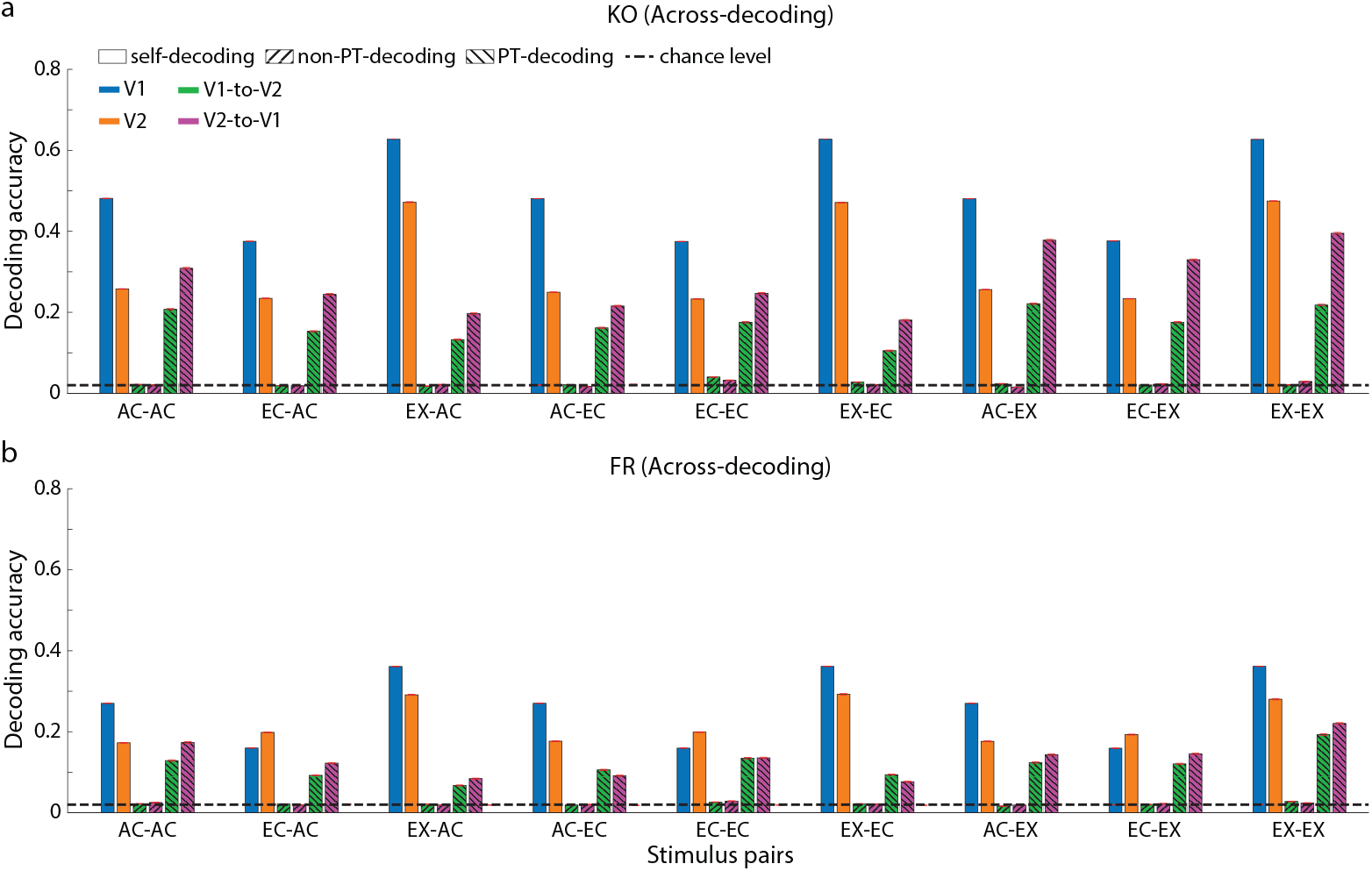
Across-region population decoding for all rendering pairs. (a) Across-region decoding accuracies for monkey KO. (see caption of Fig.4 for definition). The x-axis indicates the specific train-test rendering pair (e.g., ‘AC-EC’ means train on AC, test on EC). For each pair, four conditions are shown by color: V1 self-decoding (blue), V2 self-decoding (orange), V1-to-V2 decoding (green), and V2-to-V1 decoding (magenta). Within each colored group, different bar styles represent the decoding type as defined in the legend (self-decoding, non-PT-decoding, PT-decoding). The dashed line indicates the 2% chance level. Error bars show SEM. (b) Across-region decoding accuracies for monkey FR. Panel details are as in (a).

### Invariant Geometric Structures in Models of the Ventral Visual Stream

To further understand the importance of the geometric structure in a population code, we compared our neural data with models of the ventral visual stream. We experimented on two widely used Convolutional Neural Networks (CNNs): AlexNet (Krizhevsky et al., 2017) and VGG (Simonyan and Zisserman, 2015). These two models have been shown to encode visual information with a relatively high degree of representational similarity to regions of the ventral visual system, particularly V1 and V2 (Laskar et al., 2020; Schrimpf et al., 2018).

First, we measured the single-unit tuning correlations in both models (Fig.2.f; see Methods). We found that units in the first two pooling layers of AlexNet, unit tuning correlations were well above chance level for all rendering pairs. Similar to the biological neurons, units from the pooling layer 1 had higher tuning correlations than those in the pooling layer 2. This was also true for the pooling layer 3, although tuning correlations only reached significance for one-pair rendering sets. A similar hierarchical pattern was found for VGG, but only for units beyond the pooling layer 3, possibly due to the smaller receptive fields in its earlier pooling layers. Overall, a similar proportion of units from the models displayed cue-invariance properties, comparable to those found in V1 and V2. However, this proportion generally decreased along the successive layers of the models.

At the population level, we performed RSA and the cue-transfer decoding. RSA revealed that model unit populations had a similar level of across-rendering representational similarity pairs to the neural data (Fig.9.b). In VGG, the pooling layer 3 showed increased representational similarity, which may also be explained by its larger receptive fields compared to those in preceding layers. For cue-transfer decoding, we found that without geometric alignment, all layers of AlexNet and VGG exhibited low but above-chance accuracy, similar to the neural data (Fig.3.d). With the Procrustes Transformation, however, cue-transfer decoding accuracies in both models closely approached the self-decoding upper bounds (for AlexNet, PT accuracy was, on average, 75.26% of the self-decoding level [95% CI: 74.92%, 75.62%] in Pool 1, 89.77% [95% CI: 89.63%, 89.92%] in Pool 2, and 67.16% [95% CI: 66.81%, 67.50%] in Pool 5; for VGG, the corresponding values were 22.71% [95% CI: 22.54%, 22.88%] in Pool 1, 59.04% [95% CI: 58.78%, 59.29%] in Pool 2, 66.51% [95% CI: 66.30%, 66.72%] in Pool 3, 63.11% [95% CI: 62.93%, 63.29%] in Pool 4, and 52.05% [95% CI: 51.68%, 52.43%] in Pool 5). Hence, similarly to the neuronal data, the geometric structure of model unit populations displayed robust cue-invariant properties.

### Effect of Population Size on Cue-Invariant Properties of Biological Neurons and Model Units

To investigate how population size contributes to the geometric cue-invariant properties, we estimated cue-transfer decoding accuracies for progressively larger groups of neurons (see Methods). In all cue-transfer decoding scenarios, increasing the population size improved both self-decoding and PT-coding accuracy (Fig.4.c). These results suggest that the geometric structure of the population code becomes more stable and robust as the population size increases. The corresponding increase in within-decoding accuracy (Fig.13) also supports this.

**Figure 13:**
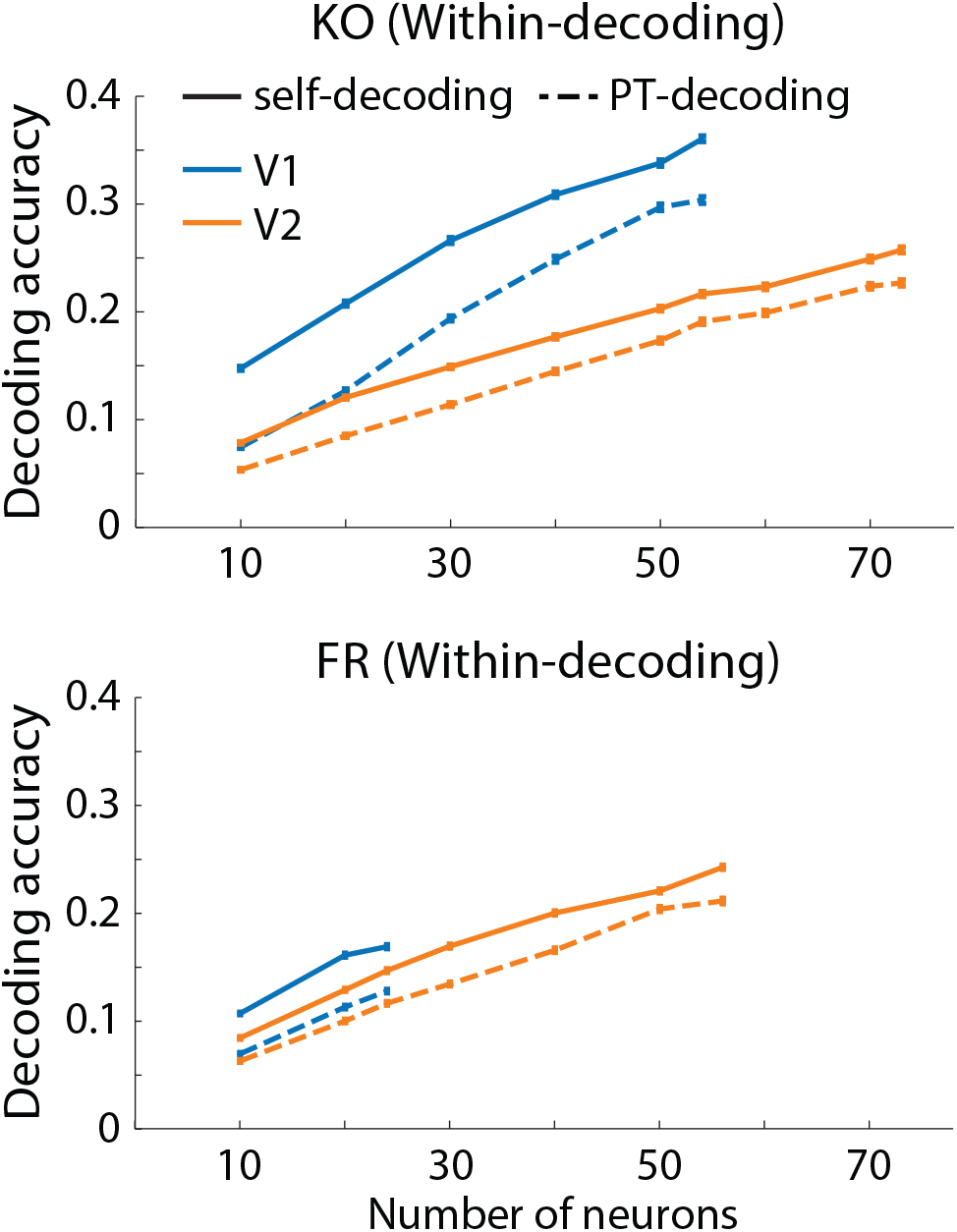
Effect of population size on ‘within-decoding’ accuracy. ‘Within-decoding’ accuracy for V1 (blue) and V2 (orange) as a function of the number of neurons, for monkey KO (top) and FR (bottom). Solid and dashed lines represent self-decoding and PT-decoding accuracy, respectively. The small horizontal bars centered on each data point represent the SEM.

However, the rate of improvement in transfer decoding accuracy tended to slow down with increasing population size. While our recorded neural populations were limited in size, the large number of units in the CNN models allowed us to estimate the population size at which performance saturates. In the CNNs, the cue-transfer decoding accuracy reached a plateau with population sizes between 250 and 300 units (Fig.4.d). This suggests that with a sufficiently large population, the geometric structure of the neural codes in V1 and V2 would also become highly stable. We conclude that, given that cortical hypercolumns in areas like V1 are highly overcomplete, it is plausible that the underlying geometric structure for boundary concepts is indeed cue-invariant in the brain.

### Geometric Invariance without Tuning Correlation in Simple and Complex Cell Models

Finally, we tested whether classical models of simple and complex cells could account for the single-neuron tuning correlations observed in V1 and V2 (Heitger et al., 1992; Hubel and Wiesel, 1968). We used Gabor filter-based models to simulate neural responses to the same set of stimuli and measure their tuning correlations (Fig.14). The results show that the sparse responses of these model cells could not explain the degree of cue-invariance measured in individual V1 and V2 neurons. We then examined whether these models could replicate the population-level cue-transfer decoding results. We measured cue-transfer decoding accuracies using the population responses of simple and complex cell models (see Methods). Self-decoding accuracies of the models were close to optimal, and just as with the neural data, non-PT cue-transfer decoding was at chance while PT cue-transfer decoding was significantly above it (Fig.15.a). Using all model neurons, simple cells approximately doubled PT cue-transfer decoding, indicating an effect of population size (Fig.15.b). When the model population size was comparable to the number of recorded neurons, the PT decoding accuracies were also on par with the neural data. Notably, while the Gabor models achieved near-optimal self-decoding, their PT decoding performance was not substantially higher than that of the neural data, which operates with much lower self-decoding accuracy. This demonstrates that classical Gabor models are sufficient to produce a cue-invariant geometric structure, though perhaps not with the same efficiency relative to their discriminability as the brain.

**Figure 14:**
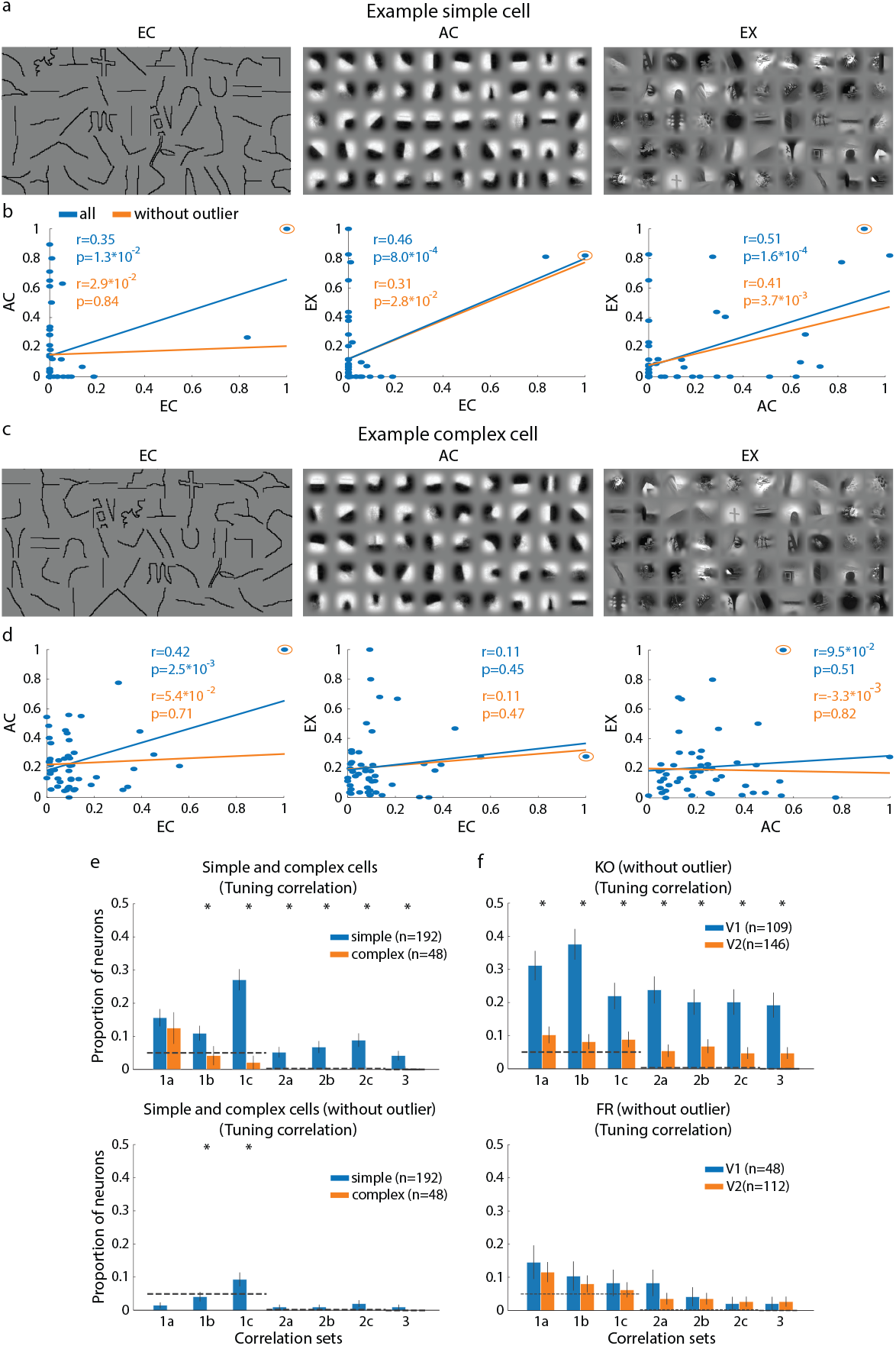
Tuning correlation of simple and complex cell models. (a, b) Tuning correlation analysis of an example simple cell. (a) Rank-ordered stimuli of each rendering based on the response of the example simple cell. (b) Scatter plots of the cell’s responses for each rendering pair. The blue line indicates the correlation using all stimuli. The orange line indicates the correlation after removing the single “outlier” stimulus (circled orange point) that most strongly drives the correlation. Insets display the Pearson correlation and p-value before (blue) and after (orange) outlier removal. (c,d) Tuning correlation analysis of an example complex cell. Panel details are as in (a, b). (e) Proportion of model cells with significant tuning correlations. Results are shown for simple (blue) and complex (orange) cells, both for the full dataset (top) and after removing the outlier stimulus for each cell (bottom). The bar chart format is the same as in Fig.2d. (f) Proportion of biological neurons with significant tuning correlations after outlier removal. Results for V1 and V2 neurons from monkey KO (top) and FR (bottom) are shown for comparison. The bar chart format is the same as in Fig.2d.

## Discussion

This study makes four key contributions to the understanding of cue-invariant neural representation in the visual cortex. First, we examined neural responses in macaque V1 and V2 to local surface boundary elements that capture the variability and complexity of shapes in natural scenes, extending the study of visual coding beyond orientation tuning. We tested responses to three renderings of diverse boundary patterns—pure contours, simplified cartoons, and naturalistic photographs. This allowed us to assess how neural representations change as the stimuli transition from abstract outlines to forms embedded in more complex images. Second, laminar recordings in both V1 and V2 enabled a comparative analysis across cortical layers. A substantial proportion of neurons in both areas exhibited significant two-way and three-way tuning correlations across the different renderings. At the single-neuron level, V1 neurons showed a modest but significant correlation of tuning curves. Our layer-wise analysis revealed that cue-invariance at the single-neuron level unexpectedly decreases along the ventral visual stream. Third, we developed a cue-transfer decoding method to assess cue invariance at the population level. This revealed that local boundary patterns share similar geometric structures in their neural manifolds across renderings, differing primarily by rotation, scaling, and translation. Thus, cue-invariance is not embedded in raw population responses but in the geometry of the population code. This geometric structure is preserved across different neuronal subpopulations within each area, and even between V1 and V2, suggesting that cue-invariant representation resides in population geometry and survives transformations along the visual hierarchy that alter tuning specificity and invariance at the single-neuron level. Fourth, we found strikingly similar cue-invariant geometric structures in deep neural network models of the ventral visual stream. In models such as AlexNet and VGG, given a sufficiently large neuron pool, this geometry is almost perfectly preserved across most layers, with some degradation only at the final pooling stages. This indicates that early and intermediate layers—analogous to early and intermediate visual areas—contain enough information for cue-transfer decoding. Such geometric invariance may underlie both the cue-invariant object recognition capacity of deep networks and biological visual systems.

### Cue-invariance Properties of Individual V1 and V2 neurons

Cue-invariance in individual neurons is typically assessed by determining whether a neuron exhibits the same tuning curve for a given perceptual attribute, such as bar orientation, when that attribute is defined by different visual cues. Classic work by Hubel and Wiesel (Hubel and Wiesel, 1968) examined orientation selectivity in V1 using both bars and edges, assuming that neurons functioned as cue-invariant edge detectors. Similarly, Connor et al. (Connor et al., 2007) reported that V4 neurons tuned to junction parameters maintained their selectivity regardless of whether junctions were defined by contours or contrast edges. Other studies have extended this approach to higher-level perceptual constructs such as figure-ground segregation, examining invariance across texture, luminance, stereo, and motion cues (Zipser et al., 1996). These investigations share the underlying assumption that a neuron encoding an abstract representation should respond similarly to that representation regardless of the visual cue defining it.

Our study extends the analysis of cue-invariance in two ways. First, rather than varying a single parametric attribute, we tested individual-neuron tuning to a diverse set of local boundary patterns derived from natural scenes, presented in three renderings: contours, cartoons, and natural photographs. This approach allows us to probe invariance across richer and more ecologically valid stimulus variations. Second, we compared tuning correlations not only between contour and cartoon renderings, but also with their natural photographic counterparts.

We quantified the fraction of neurons showing significant above-chance tuning correlations for one, two, or all three cue pairs (Fig.2). Unexpectedly, cue-invariance at the single-neuron level tended to decrease from V1 to V2, despite the ventral stream’s well-established increase in invariance to transformations such as translation, rotation, and scaling. For example, tuning correlation between EC and EX was strongest in V1 Layer 4, and correlations across all cue pairs were generally higher in V1 than in V2. This counterintuitive pattern likely reflects a trade-off: V1 neurons have highly specific spatial and feature tuning within their local receptive fields, whereas the increase in geometric invariance at higher stages can reduce discriminability for this particular set of fine-scale boundary features, thereby lowering cue-invariance in tuning curves.

A similar trade-off emerges in CNNs (Fig.2.f). Cue-invariance for these boundary features dropped beyond Pool 1 in AlexNet and beyond Pool 3 in VGG. Because our cue-invariance measure is sensitive to spatial and feature precision—a bar shifted in position is considered a different bar—any increase in translation or rotation robustness can reduce measured cue-invariance. Consistently, Gabor-based simple-cell models exhibited stronger cue-invariance for these stimuli than complex-cell models (Fig.15.a), supporting the idea that reduced spatial specificity in complex cells diminishes cross-cue tuning correlations.

**Figure 15:**
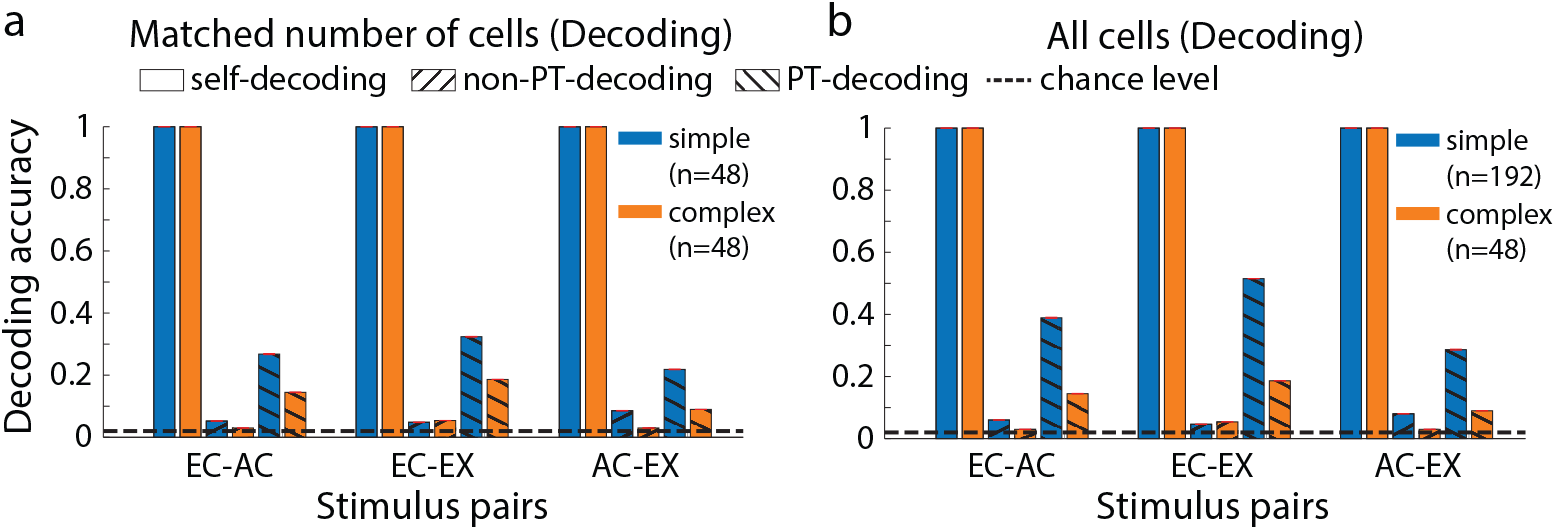
Population decoding for simple and complex cell models. (a) Population decoding accuracy for model simple (blue) and complex (orange) cells with matched population sizes. The simple cell model (originally n=192) was down-sampled to n=48 to match the size of the complex cell model population. For each model type, three conditions are shown: self-decoding (plain bars), non-PT-decoding (forward-slash hatch), and PT-decoding (backslash). The dashed line indicates the 2% chance level. (b) Population decoding accuracy using the full model populations (n=192 for simple cells and n=48 for complex cells). Panel details are as in (a).

Consistent with the observation that V1 layer 4 neurons at the individual level exhibited a stronger cue-invariant tuning correlation than V1 superficial layers and V2 neurons, we found that Gabor-based simple cells also exhibit a stronger degree of cue-invariance than complex cell models. Nevertheless, V1 and V2 neurons display a finer sensitivity and discriminability than Gabor filters. They responded in a more graded fashion across the entire stimulus set and maintained cue-invariant correlations even when the optimal stimulus (with the highest geometric mean response across two cues) was removed (Fig.14.e)—a robustness not seen in Gabor models. This might suggest that the neuron’s feature selectivity might be more nuanced and complex than an oriented Gabor-based model. Recent large two-photon calcium imaging in the awake monkey’s V1 (Tang et al., 2018) revealed that individual neurons’ selectivity is more diverse and complex, resulting in finer discrimination among these boundary patterns.

### Cue-Invariance in the Geometry of the Neuronal Population Codes in V1 and V2

Although a subset of V1 and V2 neurons exhibited significant cross-cue tuning correlations at the single-neuron level, these correlations were modest, and their contribution to the brain’s invariant representation of boundary concepts remains unclear. We therefore considered the possibility that cue-invariant representations arise not from the invariant tuning of individual neurons, but from the geometric structure of distributed population codes (Chung and Abbott, 2021; Kriegeskorte and Wei, 2021). One way to characterize this structure is by assessing whether the relative distances between neural responses to different boundary concepts are preserved across rendering cues. To test this, we first applied representational similarity analysis (RSA) (Kriegeskorte et al., 2008), which revealed significant cross-cue similarity in pairwise distances between neural representations of boundary concepts. However, RSA is limited to distance-based comparisons and does not capture the full geometric transformation between population manifolds.

To address this, we applied a cue-transfer decoding approach to assess the alignment of neural manifolds across rendering cues. We found that these manifolds are related by geometric transformations—including translation, scaling, and, importantly, rotation. While centering and normalization reduced differences due to physical factors such as contrast, they were insufficient for full cross-cue generalization. By contrast, applying a Procrustes Transformation markedly improved cue-transfer decoding performance, indicating that the geometry of boundary-concept manifolds is preserved across different renderings and can be related by geometric transformations. Notably, this stability does not depend on cue-invariant tuning at the level of individual neurons. Cue-transfer decoding remained robust under neuron-shuffle controls: when tuning preferences were randomly reassigned across neurons within each cue (Fig.3.c;Fig.11.a), decoding accuracy stayed high. The same geometric structure was preserved not only between independent populations within an area, but also between populations across V1 and V2 (Fig.4.a-b). Crucially, this structure is not trivially imposed: when population codes were shuffled across stimuli or the stimulus–response matrix was randomly permuted (Fig.11.b-c), cue-transfer decoding dropped to chance. Thus, the observed geometric alignment is both genuine and non-trivial.

The stability of manifold geometry appears stronger in V1 than in V2, possibly because V2 incorporates greater invariance to local position, scale, or rotation in stimuli. This added invariance may reduce the discriminability of fine-grained boundary distinctions, leading to a blurring of geometric relationships among certain boundary concepts. As object representations acquire increasing transformation invariance in higher visual areas (e.g., V4, IT), the precision of boundary concepts may diminish, reflecting a trade-off between specificity and generalization. Such a trade-off points to a division of computational labor: early visual areas like V1 and V2—equipped with precise spatial and orientation tuning, as well as recurrent connections implicated in boundary processing (Fitzpatrick, 1996; Gilbert et al., 2000; Lee and Mumford, 2003)—may be particularly well suited to encode cue-invariant boundary concepts, not through invariant tuning of single neurons, but through stable manifold geometry at the level of population activity.

The preservation of manifold geometry across cues, as assessed by cue-transfer decoding in our neurophysiological data, is, however, not complete. Indeed, the neurons we analyzed represent only a minute fraction of the hundreds of millions present in macaque V1 and V2. Additionally, our data, concatenated across multiple sessions, limits the evaluation of the beneficial or detrimental effect of neural correlations on the neuronal population information and cue-invariance properties (Abbott and Dayan, 1999; Farzmahdi et al., 2025; Kohn et al., 2016). Still, we found that manifold preservation systematically increases as more neurons are incorporated (Fig.4.c; Fig.13), indicating that the stability of the population geometry improves with population size. Remarkably, similar geometric invariance was also observed in CNNs trained on object classification (e.g., AlexNet, VGG; Fig.3.d), suggesting that both macaque visual cortex and CNNs may rely on comparable strategies for cue-invariant abstraction by embedding boundary concepts into manifolds that preserve geometry across cues. This parallel provides a useful framework for studying how the stability of geometric structure depends on the number of sampled neurons. Indeed, cue-transfer decoding applied to progressively larger sets of units in early layers showed that invariance approaches near perfection (Fig.4.d). With sufficiently large populations, the manifold geometry is effectively preserved, offering a robust and scalable coding strategy for abstraction and cue-invariant representation.

Finally, the observed scaling and translation of boundary-concept manifolds across rendering cues likely reflect differences in contrast sensitivity and individual neuronal feature preferences. The rationale for why these manifolds undergo rotation across cues is less direct. One possibility is that rotation provides representational advantages, such as reducing interference between overlapping neural codes—a mechanism proposed in the context of memory separation and contextual encoding (Libby and Buschman, 2021). Another possibility is that rotated geometry helps preserve perceptual distinctiveness across cues. Although line drawings, cartoons, and photographs can all support similar interpretations, they remain perceptually distinct. Rotation in representational geometry may therefore promote perceptual segregation while still enabling downstream neurons to construct explicit representations of abstract concepts. Because manifold geometries across cues are related by a geometric transformation, this construction should be straightforward. Scaling and translation are already known to be implemented in the brain through canonical mechanisms such as divisive normalization and gain control, and recent theoretical work suggests that population-level rotations are biologically plausible through circuit dynamics (Pu et al., 2024; Rule and O’Leary, 2022; Rule et al., 2020). Thus, by relying on population activity rather than cue-invariant tuning of single neurons, the visual system can maintain perceptual distinctions among concepts rendered in different cues while still achieving stable, abstract, and semantic representations through the geometry of population codes.

## Materials and Methods

### Animal Preparation

Two adult male Rhesus macaque monkeys (Macaca mulatta) participated in the study (monkey KO and FR). Under general anesthesia, each animal was implanted with a head fixation post, scleral search coils, and a recording chamber overlying the occipital operculum, with its center roughly at the border between areas V2 and V1 (more details can be found in (Smith et al., 2007)). All procedures were approved by the Institutional Animal Care and Use Committee of Carnegie Mellon University and complied with the guidelines set forth in the United States Public Health Service Guide for the Care and Use of Laboratory Animals.

### Visual Stimuli

This study aimed to analyze the responses of V1 and V2 neurons to a set of boundary surface stimuli with varying shapes that reflect the natural statistics of the visual environment. First, we extracted a large number of patches from 2D natural scenes in the Berkeley Segmentation Dataset (Chen et al., 2013; Martin et al., 2001). We used edge annotations from human experts and the associated edge segmentation binary maps from all images to collect a total of 5818 patches of size 33×33 and 45×45 pixels (Fig.1.a). Then, we computed pairwise shape distances between the segmented edges of each patch using the Hausdorff shape metric. We clustered the whole set of edges using affinity propagation (Frey and Dueck, 2007) into 50 clusters. We found that this number of clusters was sufficient to approximate the variety of surface boundary shapes occurring in the natural scenes of the database. Advantageously, this example-based clustering method generates clusters whose centers are actual examples from the dataset. These centers formed the set of ‘Edge Contours’ (EC) stimuli (Fig.5.a). The original patches extracted from the natural scenes, from which the EC stimuli originated, formed the set of ‘Example’ (EX) stimuli (Fig.5.b). Finally, to model the patch appearance, we performed a principal components analysis (PCA) on the aligned and normalized edge patches of each cluster. The first component was visually similar (albeit with possible reversal of edge luminance polarity) to the mean luminance patch of the cluster. The mean patches of each cluster formed the set of ‘Appearance Contour’ (AC) stimuli (Fig.5.c). To prevent border artifacts when measuring individual neurons’ responses, a circular smoothing aperture was applied to the patch to remove any abrupt edge information (Fig.5.d). Two stimulus sizes (33×33 and 45×45 pixels) were used to match the average receptive field sizes of V1 and V2 neurons (the field of view is 1^°^ for V1 and 1.5^°^ for V2). The three renderings of the selected 50 surface boundary contours resulted in a dataset of 150 stimuli used for the visual fixation task (Fig.1.c).

### Visual Fixation Task

Eye movements were recorded using a scleral search coil system (Riverbend Instruments, Birmingham, AL), sampled at 200 Hz. Behavioral task control was managed using NIMH Cortex. After recovery from surgery, each animal was trained to perform a standard fixation task. A trial began with the appearance of a central fixation dot. The animal was required to fixate on the dot within 3000 ms and then maintain its eye position within a 0.8° window of the central point for 200-350 ms. At the end of this period, a visual stimulus appeared at a specific location on the screen (see below Receptive Field Mapping), and the animal was required to maintain its fixation for at least 300 ms. If all these conditions were met, the trial was counted as successful, and the animal received a liquid reward. Otherwise, the trial was aborted. Stimuli were presented in blocks, with each block containing only one rendering type. Within a block, each stimulus was presented 10 times in a randomized, non-consecutive order.

### Neurophysiological Recordings

We used a 2×12-channel laminar probe (Alpha-Omega, Alpharetta, USA; 150 *µ*m inter-contact distance; 300 *µ*m diameter; 1 MΩ impedance for each channel) to record neural activity in cortical areas V1 and V2. The probe was inserted through the dura into the V1 operculum using a hydraulic Microdrive (Narishige, Tokyo, Japan). V1 neural activity was identified as the activity first detected upon the insertion and lowering of the probe. V2 neural activity was recorded by increasing the depth of the probe. In each region, the probe was positioned to ensure that at least one contact had a well-isolated spike waveform (the reference neuron) and that activity from the same region (V1 or V2) was recorded on as many of the remaining contacts as possible. The discrimination between V1 and V2 activity was assessed based on a clear decrease in the neural activity (due to the presence of white matter) and on the size of the receptive field of the reference contact (see below Receptive Field Mapping). Neurophysiological signals were recorded using the Blackrock acquisition system (Blackrock, Salt Lake City, USA). Data recording was synchronized with the beginning of the trial, and the timing of all trial events was recorded simultaneously with the neural activity. For each channel, the raw activity was parsed into spiking activity (high pass filter at 250 Hz and threshold at 3.5 times the root mean square of the activity) and Local Field Potentials (LFP, low pass filter at 250 Hz). Spiking waveforms were sorted offline using a competitive mixture decomposition method and manually refined based on their shape and the interspike interval using custom time amplitude window discrimination software written in MATLAB (Kelly et al., 2007) (available at https://github.com/smithlabvision/spikesort). Each session typically consisted of a single electrode penetration.

### Receptive Field Mapping

During each experimental session, the receptive field of the reference neuron (see above Neurophysiological Recordings) was determined using a hand-mapping bar stimulus while the animal maintained fixation. Subsequent visual stimuli were displayed at the center of this position. During offline analysis, we refined these parameters using the minimum response field technique with a small oriented bar to map the receptive fields of all the neurons across all contacts. Receptive field positions ranged from 2 to 5^°^ eccentricity in the lower visual field, and diameters ranged from 1 to 1.5^°^.

### Laminar Analysis and Early/Late Responses

For each penetration, channels were divided into Superficial Layers, Layer 4, and Deep Layers (Fig.6.a). To do so, we used current-source density analysis (CSD) and the onset latency of the neural activity to estimate the location of Layer 4, corresponding to lateral geniculate nuclei or cortical input activity. A typical CSD plot of the LFP activity of V1 and V2 cortical regions displays a strong source of input currents whose location and width were used to estimate the location of Layer 4 (Pettersen et al., 2006) (csdplotter toolbox (https://github.com/espenhgn/CSDplotter, Fig.6.b). This estimation was independently verified using onset latency, confirming Layer 4 as the one with the shortest latency across contacts (Maier et al., 2010) (Fig.6.c). Early (40-110 ms) and late (110-180 ms) responses were defined based on the average peristimulus time histogram (PSTH) across the responses of all neurons within each layer (Fig.6.d).

### Single Neuron Tuning Curves and Tuning Correlation

To measure the tuning selectivity of single neurons, we computed the averaged response of each neuron to all stimuli across the 10 repeated trials. For each neuron, three ‘tuning curves’ were obtained by rank-ordering their responses for each stimulus, grouped into the three renderings (EC, EX, and AC). Single neuron’s ‘tuning correlation’ was measured using the Pearson correlation between pairs of tuning curves (*corr* function in MATLAB). We assessed the significance of the tuning correlation based on the p-value of the Pearson correlation (significance level *p <* 0.05). To estimate how single neuron cue-invariance characteristics were distributed over the whole population of neurons, we computed the proportion of neurons that exhibit significant positive tuning correlation for different correlation sets. Correlation sets are: EC-AC (1a); EC-EX (1b); AC-EX (1c); EC-AC and EC-EX (2a); EC-AC and AC-EX (2b); EC-EX and AC-EX (2c); EC-AC and AC-EX and EC-EX (3). A neuron was included in the proportion of a given correlation set if its tuning correlations for all the pairs of renderings included in the set were significant and positive. The same neuron could be counted in multiple correlation sets. For example, a neuron with significant positive tuning correlations for both ‘EC-AC’ and ‘EC-EX’ correlation sets would be counted in the one-pair sets (1a and 1b) and also in the two-pair set (2a). Standard errors of the proportion of neurons for each correlation set were computed using the standard error of sample proportion (SSE) (Brightwell and Dransfield, 2013):

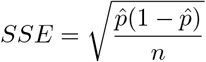

where 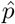 is the proportion of neurons and *n* is the number of neurons (sample size).

To measure the significance of the proportion of neurons for each correlation set, a proportion confidence interval was measured using the Wilson score interval (Wilson, 1927):

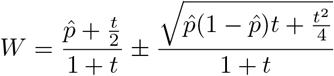

where *W* is the Wilson interval with upper and lower bounds. *t* is a random variable that indicates the difference between the true proportion and the observed proportion, which follows a standard normal distribution. 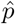 is the observed proportion of significant neurons. Here we use a *t* = 1.96 that corresponds to a confidence level of 95%.

A proportion of neurons was significant if the lower bound of the Wilson score interval was above the chance level. The chance level of the proportions for correlation sets with one-pair rendering sets was 5% (significance level of the Pearson correlation). The chance levels of correlation sets with two- and three-pair rendering sets were the square and cube of the chance level of one-pair rendering sets (2.5 × 10^−3^, and 1.25 × 10^−4^ for two- and three-pair rendering sets, respectively).

To compare the proportion of neurons between V1 and V2 regions, a Comparison of Population Proportion (CPP) was conducted (Brightwell and Dransfield, 2013). Given 2 proportions of neurons *p*_1_ and *p*_2_, the test statistic for testing the null hypothesis *H*_0_ : *p*_1_ − *p*_2_ = 0 is:

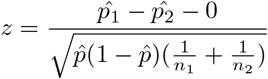

with:

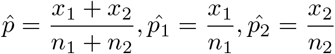

where *z* should follow a standard normal distribution; *x*_1_ and *x*_2_ are the number of neurons that showed a significant positive correlation in the two proportions of neurons; *n*_1_ and *n*_2_ are the total number of neurons.

### Representational Similarity Analysis

We used representational similarity analysis (RSA) as a measure of the correlation between 2 population responses to a rendering pair (Kriegeskorte et al., 2008). First, we computed the matrix representing the similarity of the population responses for each rendering (RSA matrix). The RSA matrix for each type of stimulus is calculated as the Pearson correlation matrix between the population tuning and itself. For each rendering, we calculated the pair-wise Pearson correlation (*corr* function in MATLAB) between the population response vectors to all pairs of stimuli (1225 pairs for 50 stimuli). RSA was estimated for each rendering pair within the one-pair rendering set (i.e., ‘EC-AC’, ‘EC-EX’, and ‘AC-EX’).

The confidence intervals were obtained by bootstrapping the set of trials used to compute the population responses (1000 bootstrap samples). The significance of the representation similarity correlations was assessed by the exclusion of the chance level (0%) from the 95% confidence intervals.

### Procrustes Transformation

Procrustes Transformation (PT) aims to align two shapes, each composed of landmark points, by finding the optimal transformations that map one to the other. In our analysis, we define two geometric shapes: a ‘target’ and a ‘comparison’. Each shape is composed of landmark points, where a single landmark represents the population response vector (the activity of all neurons) to one stimulus. Together, these landmarks map out the structure of the neural responses in a high-dimensional space. To align the target and comparison shapes, PT seeks to minimize the differences between the landmark points of both shapes using rotations, reflections, scaling, and translations (Gower, 1975). The Procrustes Transformation is expressed as:

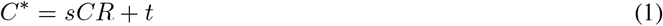

where *C* is an *n*-by-*p* matrix representing the comparison shape undergoing the Procrustes transformation. Here, *n* represents the number of landmark points (one for each of the 50 stimuli), while *p* is the number of dimensions (neurons). Each row of *C* thus corresponds to a *p*-dimensional landmark point. The transformed comparison shape is represented by the *n*-by-*p* matrix *C*^∗^. The transformation itself is defined by *R*, a *p*-by-*p* matrix for rotations and reflections; *s*, a scalar for scaling; and *t*, an *n*-by-*p* matrix for translation.

The Procrustes Transformation aims to optimize *s, R*, and *t* so that the sum of the squared differences (Euclidean distances) between corresponding landmark points of the target and transformed comparison shapes is minimized. The residual of the sum of the squared differences after transformation, *S*, is given by:

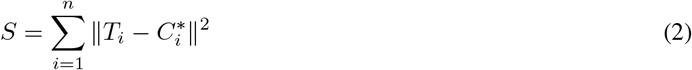

where *T*_*i*_ and 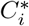 are the *i*-th landmark point of the target and transformed comparison shapes, respectively; *n* is the number of landmark points.

### Cue-Transfer Decoding

To obtain a measure of cue-invariance processing across renderings in the ventral visual stream, we developed a “cue-transfer” decoding analysis to measure how accurately a decoder trained on one visual cue (one stimulus rendering) could decode a second cue (a different stimulus rendering). We used support vector machines (SVM) to measure the cue-transfer decoding accuracy of the 50 stimuli between two renderings, with and without a geometric alignment of the neural population code via Procrustes Transformation. Specifically, PT was used to align the neural population’s responses to stimuli that were presented in two different rendering conditions: ‘rendering 1’ (the target shape) and ‘rendering 2’ (the comparison shape), as shown in Fig.10.a.

To prepare the neural data for the PT analysis, spike counts were first calculated in a 300-ms window from 30 ms to 330 ms after stimulus onset. We then applied z-scoring to each neuron and then computed its mean response across all 10 trials for each of the 50 stimuli in rendering 1. The trial-averaged responses to rendering 1 corresponded to the landmarks of the target shape and were used to define the matrix *T* in equation 2 (size 50-by-*p*, where *p* is the number of neurons). To identify the optimal alignment procedure, we empirically tested different combinations of data normalization and PT. We found that using the full PT, which includes scaling and translation, resulted in slightly lower decoding accuracy (data not shown) than using rotation alone. This is likely because the scaling and translation parameters, when optimized on trial-averaged responses, can be detrimental when applied to the variability of single-trial data. Therefore, for all results presented, we adopted the combination that yielded the highest performance: applying z-scoring to the neural data, followed by a transformation consisting only of the rotational component of PT. After z-scoring, we partitioned the trials from rendering 2 into training and validation sets using a leave-one-out procedure. For each of the 50 stimuli, 9 of the 10 trials served as the training set (referred to as ‘Training trials of rendering 2’ in Fig.10.a), while the one remaining trial was held out as the validation set (‘Validation trials of rendering 2’ in Fig.10.a). First, we computed the trial-averaged responses of the training set to define the landmarks of the comparison shape (matrix *C* in equation 1). Next, PT was applied to identify the optimal transformation for mapping the comparison shape (rendering 2) onto the target shape (rendering 1), using the *procrustes* function in MATLAB. Finally, the obtained transformation was applied to the held-out validation data (the left-out trial for each stimulus). This step produced a set of transformed trials, referred to as ‘PT trials of rendering 2’ (Fig.10.a), which were now optimally aligned with the population responses of rendering 1.

An SVM classifier (using the MATLAB *fitcecoc* function) was trained on the complete set of trials (10 trials for each of the 50 stimuli) from the target rendering (‘rendering 1’ in Fig.10.b). The trained SVM was then used to predict the stimulus identity of the held-out validation data from the comparison rendering (rendering 2). Specifically, decoding was performed on both the original validation trials to yield the non-PT accuracy and on the transformed ‘PT trials’ to yield the PT accuracy (referred to as “validation trials of rendering 2” and “PT trials of rendering 2” in Fig.10.b). The previously described leave-one-out approach was crucial for preventing data leakage, as the data used for testing the decoder was never part of the set used to calculate the transformation parameters. To ensure a stable estimate of these decoding accuracies, the entire procedure was repeated across multiple trials and neuron samples. We performed 15 different trial samplings. The selection of a trial index for each stimulus was independent, which allowed us to generate more than 10 unique samples from the available data. Furthermore, to allow for a fair comparison between V1 and V2 populations of different sizes, we equalized the number of neurons by randomly down-sampling the larger population 50 times. In total, the final reported accuracy for any given condition reflects an average over 750 repetitions (15 trial samplings × 50 neuron samplings). Finally, for each pair of renderings, the analysis was performed reciprocally. For example, for the AC-EC pair, one procedure involved training on AC and decoding on EC, and the reverse was also performed. The final reported decoding accuracy for the rendering pair is the average of these two estimations.

### Upper and Lower Bounds on Cue-Transfer Decoding

We established an upper bound on decoding performance, termed ‘self-decoding’ (Fig.10.c), by training and testing an SVM on responses to the same rendering. The self-decoding accuracy was estimated using a 10-fold cross-validation (implemented with MATLAB’s *crossval* and *kfoldLoss* functions). To estimate a lower bound, we retained the general PT decoding framework (Fig.10.a and b) but shuffled the correspondence between landmarks from different renderings (‘shuffle of stimulus labels’ in Fig.11.a). Specifically, after obtaining the 50-by-*p* landmark matrices for the target and comparison shapes, we shuffled the rows of the comparison matrix, rearranging the sequence of its 50 landmarks. This process destroyed the one-to-one correspondence between the landmarks in the two matrices. In other words, PT was forced to find a transformation between a randomly perturbed comparison shape and the target shape. By eliminating the meaningful correspondences, any decoding accuracy achieved under this condition reflected chance performance, thus providing a lower bound accuracy for the cue-transfer PT decoding. The decoding accuracy for this shuffled data approached the theoretical chance level of 2% for 50 classes (Fig.11.b,c).

### Effect of Single-Neuron Tuning Correlations on Cue-Transfer Decoding

To measure the effect of single-neuron tuning correlations on cue-transfer decoding, we conducted a neuron-shuffling experiment (‘shuffle of neuron indices’ in Fig.11.a). In this control, we removed the neuron-by-neuron correspondence between the two sets of population responses, i.e., the same population’s responses to the target and comparison renderings. Specifically, the procedure randomized the neuron indices within the neural response to the comparison rendering. By severing the one-to-one correspondence of individual neurons across the two response sets, this shuffle effectively removes any influence from their specific tuning correlations on the decoding outcome.

### Within and Across Region Transfer Decoding

We performed two additional analyses to test the stability of the population geometry within and across cortical areas. For the ‘within-decoding’ analysis, we tested whether the geometric structure of the neural code was consistent across different subsets of neurons in the same cortical area. To do this, the full population of neurons within one area (e.g., V1) was randomly split into two subpopulations of equal size. We then performed a transfer decoding analysis between them, calculating both non-PT and PT accuracy. This procedure was analogous to the main cue-transfer analysis in Fig.10.b, with one key difference: the ‘target’ and ‘comparison’ population codes were drawn from the two different subpopulations, but for the same stimulus rendering, rather than from the same population responding to different renderings.

For the ‘across-decoding’ analysis, we tested if this geometric structure was shared between V1 and V2. We applied cue-transfer decoding between the V1 and V2 populations for stimuli of either the same or different renderings. Decoders were trained on the V1 population to decode V2 responses (‘V1-to-V2 decoding’) and vice versa (‘V2-to-V1 decoding’). To ensure a fair comparison, the larger V2 population was randomly down-sampled to match the number of neurons in the V1 population for these analyses.

### Neural Network Models of the Ventral Visual Stream

AlexNet (Krizhevsky et al., 2017) is one of the early era deep convolution neural networks (CNN) designed for ImageNet image classification tasks. It is composed of 5 convolution layers and 3 fully connected layers. VGG (Simonyan and Zisserman, 2015) is another signature deep CNN that aims to reduce the number of parameters, resulting in more efficient training times. VGG-19 has 16 convolution layers grouped into 5 blocks and 3 fully connected layers.

For our neural network analyses, we used the MATLAB implementations of AlexNet and VGG-19. For each model, we extracted the activations from the units (“channels”) at the center of the feature map of each pooling layer. To prepare the input, our stimulus images were first rescaled to match the receptive field size of the recorded neurons and then zero-padded to fit the required input dimensions of the networks. The activation level of each selected channel was then treated as the firing rate of an artificial neuron for all subsequent analyses.

To investigate whether the cue-invariant structure of the neuronal data was also present in computational models, we applied our cue-transfer decoding analysis to the intermediate pooling layers of the CNNs. For each layer, we passed our stimulus set through the network to generate a 50-by-*p* matrix of activation responses, where each row corresponded to a stimulus and each column to a unit. This output matrix was subsequently treated as the mean response of the layer to the stimulus set. To simulate the trial-by-trial variability analogous to that in neuronal data, we then generated 10 trials for each stimulus by drawing random numbers from Poisson distributions centered on these mean responses. By formatting the CNN data to mimic the structure of the neuronal recordings, we were able to apply the identical cue-transfer decoding scheme (Fig.10) to the network’s responses, yielding the results presented in Fig.3d.

### Effect of Population Size on Cue-Transfer Decoding

To measure the effect of population size on cue-transfer decoding accuracy, we recalculated our decoding analyses for progressively larger groups of neurons. For each population size, we created a subset by randomly sampling neurons from the full recorded population, starting with a small subset and progressively increasing its size. This down-sampling procedure was applied to the standard cue-transfer decoding and the within-region decoding, and was also performed on the units from the artificial neural networks.

### Model of Simple and Complex Cells

The responses of simple and complex cells were modeled using Gabor filters(Granlund, 1978). Odd and even Gabor filters are defined as:

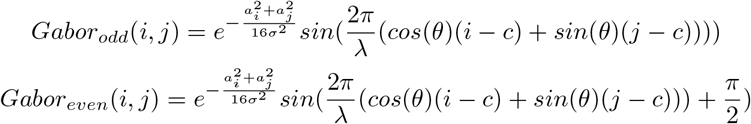

where:

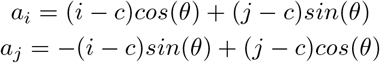

2*c* is the size of the filter in pixels, and *θ* is the orientation in radians.

To generate population responses, we created a bank of even and odd Gabor filters spanning 3 scales (20, 40, and 60 pixels) and 16 orientations (from 0° to 180° in 15° steps). Each filter in this bank was convolved with each stimulus from all three renderings. The final response for each filter was calculated as the average activation within a 3-by-3 pixel region at the center of the convolved output. This central averaging was performed to increase response variability and to simulate the potential effect of small eye position jitters around the fixation point.

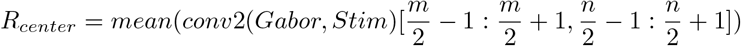

where [*m, n*] is the size of the coefficient matrix. The even and odd Gabor filters, as well as their negative counterparts, were rectified to retain only the positive coefficients. This procedure yielded a population of 192 model simple cells, derived from the combination of 3 filter scales, 16 orientations, 2 phases (even and odd), and 2 response polarities (positive and negative rectification). The response of complex cells was computed as the sum of the squared responses of the corresponding even and odd Gabor filters.

To determine if classical models of the early visual system also exhibit a cue-invariant representational structure, we applied our cue-transfer decoding analysis to the responses of the simple and complex cell models. Similar to our preprocessing of the deep neural network data, we generated simulated trial-by-trial variability by using the Gabor filter response matrices as mean rates for a Poisson random number generator, creating 10 trials for each stimulus. The cue-transfer decoding scheme, as previously described (Fig.10), was then applied to this simulated Gabor filter response data. For the simple cell analysis, we explored two variations in population size: first, using all 192 model cells, and second, down-sampling the population to 48 to match the number of complex cells. The results of these analyses are presented in Fig.15.

### Statistics

All data were analyzed using custom MATLAB code. For single neuron tuning analyses, significance was assessed by comparing the lower bounds of the confidence intervals with the chance level. Wilcoxon signed-rank tests were used for comparing independent samples. The significance level was set to *p* = 0.05 unless otherwise noted. The statistical test between two proportions described above in section ‘Single Neuron Tuning Selectivity’, assumes the difference between the two proportions follows a standard normal distribution under the null hypothesis. For the decoding analyses, the mean and standard error of the decoding accuracy were estimated using a bootstrap procedure. This involved repeatedly resampling the trials and neurons to generate a distribution of decoding accuracies, from which the final statistics were computed. SEM bars are reported on the bar charts. Comparisons between conditions (e.g., PT vs. non-PT) were performed on the full distribution of repetitions. To obtain a single summary statistic, the accuracy differences from all relevant comparisons were pooled, and the mean and a 95% confidence interval of this pooled distribution were calculated using a bootstrap procedure with 10,000 repetitions.

## Data and Code Availability

All data and custom MATLAB code used in this study have been deposited in a public GitHub repository located at: https://github.com/ZitongWang-dev/cue-invariant-population-geometry

## Acknowledgments

We thank Karen McCracken of the Center for the Neural Basis of Cognition for animal care and surgery preparations.

## Effect of Temporal Window on Single-Neuron Cue-Invariance

We also investigated whether the cue-invariant properties of individual neurons depended on the response time window by distinguishing between early (40-110 ms) and late (110-180 ms) activity. These windows may correspond to initial feedforward and later feedback processing, respectively. We re-analyzed the neural tuning invariance, distinguishing two temporal integration windows: early (40-110 ms) and late (110-180 ms) windows (see Methods, Fig.6.d). Figure 8.b,c shows the proportion of cue-invariance neurons for early and late temporal windows in V1 and V2 in both monkeys. In both monkeys, individual V1 neurons showed significant tuning correlation across all correlation pairs, whether the neural activity was analyzed within the early or the late window. The proportion of V1 neurons showing significant cue-invariance was, on average, very similar across temporal windows. For monkey KO, the proportions of V1 neurons showing significance for one-, two-, and three-pair rendering sets were 30%, 20%, and 16% in the early window, compared to 28%, 19%, and 17%in the late window. For monkey FR, the proportions were 21%, 10%, and 6% (early) versus 18%, 6%, and 4% (late). These results allow us to conclude that, in V1, the proportion of cue-invariant neurons did not depend on the window of analysis.

In V2, the results were less consistent. In the early window, the proportion of cue-invariant neurons was significantly lower in V2 than in V1 for both monkeys (CPP, *p* < 0.01; figure8.b,c). In addition, in monkey KO, the proportion of V2 neurons showing significant tuning correlation was significant across all correlation pairs. In monkey FR, this proportion was largely reduced and only reached significance for EC-EX (1b) and EC-AC and EC-EX (2a) correlation pairs. In the late window, this pattern was somewhat reversed. In monkey KO, the proportion of significant neurons was lower in V2 than in V1 (CPP, *p* < 0.01). The proportion of V2 neurons showing significant tuning correlation was largely reduced across all correlation pairs and did not reach significance for EC-EX (1b) and AC-EX (1c) correlation pairs. Furthermore, for monkey FR, the proportion of significant V2 neurons in the late window was high across all correlation sets, in stark contrast to the results from monkey KO. These contradictory findings prevent any clear conclusion about the dependence of V2 cue-invariance on the analysis window.

## Representation similarity analysis does not show invariance difference between V1 and V2

We applied representational similarity analysis (RSA) to population response patterns for each pair of renderings. In both animals, we found that representation similarity was significant for the EC-AC and EC-EX pairs in both V1 and V2 populations. For the remaining AC-EX pair, only monkey KO’s V1 population response reached statistical significance. Given that the AC and EX renderings were derived from EC, this shared origin likely explains the significant similarity in pairs that included EC, while the AC-EX pair showed no significant similarity. However, the results were not consistent across animals: V1 representational similarity was higher than V2 for the EC-AC and EC-EX pairs in monkey KO, but this difference was not significant in monkey FR. Because these RSA results were inconsistent across animals, we turned to an alternative analysis to leverage information about population structure not captured by RSA.

## Tuning correlation analysis with models of simple and complex cells

To test whether classical models of simple and complex cells could explain the tuning correlations measured in V1 and V2, we used Gabor filter-based models to simulate neural responses and measure their tuning correlations (Fig.14). We observed that the model cell responses were typically very sparse, with each cell responding preferentially to only a few stimuli. In the context of the correlation measure, we termed these highly effective stimuli “outliers” (Fig.14.a-d shows 2 examples of sparse responses of a simple and a complex cell). We computed the proportion of simple and complex cells with significant tuning correlations, both with and without these outlier stimuli (Fig.14.e). In both cases, the proportions were low or not significantly different from chance, and they were even lower after the outliers were removed. As a control, we performed the same outlier removal procedure on the biological data and found that the proportion of significant neurons was not substantially affected (Fig.14.f; Fig.2.d). These results show that classical models of simple and complex cells, with their sparse responses, do not account for the amount of cue-invariance measured in individual V1 and V2 neurons.

## References

Abbott, L. F., & Dayan, P. (1999). The effect of correlated variability on the accuracy of a population code. Neural Comput, 11(1), 91–101.

Alitto, H. J., & Usrey, W. M. (2004). Influence of contrast on orientation and temporal frequency tuning in ferret primary visual cortex. J Neurophysiol, 91(6), 2797–808.

Angelucci, A., Bijanzadeh, M., Nurminen, L., Federer, F., Merlin, S., & Bressloff, P. C. (2017). Circuits and mechanisms for surround modulation in visual cortex. Annu Rev Neurosci, 40, 425–451.

Anzai, A., Peng, X., & Van Essen, D. C. (2007). Neurons in monkey visual area v2 encode combinations of orientations. Nature Neuroscience, 10(10), 1313–1321.

Brightwell, B., & Dransfield, B. (2013). Avoid and detect statistical malpractice: Design and analysis for biologists, with r. InfluentialPoints.

Chen, L.-C., Papandreou, G., & Yuille, A. L. (2013). Learning a dictionary of shape epitomes with applications to image labeling. Proceedings of the IEEE International Conference on Computer Vision (ICCV).

Chung, S., & Abbott, L. F. (2021). Neural population geometry: An approach for understanding biological and artificial neural networks. Current Opinion in Neurobiology, 70, 137–144.

Connor, C. E., Brincat, S. L., & Pasupathy, A. (2007). Transformation of shape information in the ventral pathway. Current Opinion in Neurobiology, 17(2), 140–147.

Dobbins, A., Zucker, S. W., & Cynader, M. S. (1987). Endstopped neurons in the visual cortex as a substrate for calculating curvature. Nature, 329(6138), 438–41.

Elder, J. H. (2015). Bridging the dimensional gap: Perceptual organization of contour into two-dimensional shape. In J. Wagemans (Ed.), The oxford handbook of perceptual organization. Oxford University Press.

Elder, J. H., Oleskiw, T. D., & Fruend, I. (2018). The role of global cues in the perceptual grouping of natural shapes. Journal of Vision, 18(12), 14.

Farzmahdi, A., Kohn, A., & Coen-Cagli, R. (2025). Relating natural image statistics to patterns of response covariability in macaque primary visual cortex. Nature Communications, 16(1), 6757.

Fitzpatrick, D. (1996). The functional organization of local circuits in visual cortex: Insights from the study of tree shrew striate cortex. Cereb Cortex, 6(3), 329–41.

Freeman, J., Ziemba, C. M., Heeger, D. J., Simoncelli, E. P., & Movshon, J. A. (2013). A functional and perceptual signature of the second visual area in primates. Nature Neuroscience, 16(7), 974–981.

Frey, B. J., & Dueck, D. (2007). Clustering by passing messages between data points. Science, 315(5814), 972–976.

Geisler, W. S., & Perry, J. S. (2009). Contour statistics in natural images: Grouping across occlusions. Vis Neurosci, 26(1), 109–21.

Gilbert, C., Ito, M., Kapadia, M., & Westheimer, G. (2000). Interactions between attention, context and learning in primary visual cortex. Vision Res, 40(10-12), 1217–26.

Gower, J. C. (1975). Generalized procrustes analysis. Psychometrika, 40(1), 33–51.

Granlund, G. H. (1978). In search of a general picture processing operator. Computer Graphics and Image Processing, 8(2), 155–173.

Hegdé, J., & Van Essen, D. C. (2000). Selectivity for complex shapes in primate visual area v2. J Neurosci, 20(5), Rc61.

Hegdé, J., & Van Essen, D. C. (2007). A comparative study of shape representation in macaque visual areas v2 and v4. Cereb Cortex, 17(5), 1100–16.

Heitger, F., Rosenthaler, L., Von Der Heydt, R., Peterhans, E., & Kübler, O. (1992). Simulation of neural contour mechanisms: From simple to end-stopped cells. Vision Research, 32(5), 963–981.

Hubel, D. H., & Wiesel, T. N. (1968). Receptive fields and functional architecture of monkey striate cortex. The Journal of Physiology, 195(1), 215–243.

Kanwisher, N., & Yovel, G. (2006). The fusiform face area: A cortical region specialized for the perception of faces. Philosophical Transactions of the Royal Society B: Biological Sciences, 361(1476), 2109–2128.

Kelly, R. C., Smith, M. A., Samonds, J. M., Kohn, A., Bonds, A. B., Movshon, J. A., & Sing Lee, T. (2007). Comparison of recordings from microelectrode arrays and single electrodes in the visual cortex. The Journal of Neuroscience, 27(2), 261–264.

Kohn, A., Coen-Cagli, R., Kanitscheider, I., & Pouget, A. (2016). Correlations and neuronal population information. Annual Review of Neuroscience, 39(Volume 39, 2016), 237–256.

Kriegeskorte, N., & Kievit, R. A. (2013). Representational geometry: Integrating cognition, computation, and the brain. Trends Cogn Sci, 17(8), 401–12.

Kriegeskorte, N., & Wei, X. X. (2021). Neural tuning and representational geometry. Nat Rev Neurosci, 22(11), 703–718.

Kriegeskorte, N., Mur, M., & Bandettini, P. (2008). Representational similarity analysis - connecting the branches of systems neuroscience. Frontiers in Systems Neuroscience, 2.

Krizhevsky, A., Sutskever, I., & Hinton, G. E. (2017). Imagenet classification with deep convolutional neural networks. Commun. ACM, 60(6), 84–90.

Laskar, M. N. U., Sanchez Giraldo, L. G., & Schwartz, O. (2020). Deep neural networks capture texture sensitivity in v2. Journal of Vision, 20(7), 21–1.

Lee, T. S., & Mumford, D. (2003). Hierarchical bayesian inference in the visual cortex. J Opt Soc Am A Opt Image Sci Vis, 20(7), 1434–48.

Libby, A., & Buschman, T. J. (2021). Rotational dynamics reduce interference between sensory and memory representations. Nature Neuroscience, 24(5), 715–726.

Lieber, J. D., Oleskiw, T. D., Palmieri, L., Simoncelli, E. P., & Movshon, J. A. (2025). Responses of neurons in macaque v4 to object and texture images. bioRxiv, 2024.02.20.581273.

Liu, M., Chang, S., Chen, M., Li, P., Roe, A. W., & Hu, J. M. (2025). How shape information is coded by v4 cortical response of macaque monkey. Journal of Neurophysiology, 133(6), 2016–2028.

Maier, A., Adams, G. K., Aura, C., & Leopold, D. A. (2010). Distinct superficial and deep laminar domains of activity in the visual cortex during rest and stimulation. Frontiers in Systems Neuroscience, 4.

Martin, D., Fowlkes, C., Tal, D., & Malik, J. (2001). A database of human segmented natural images and its application to evaluating segmentation algorithms and measuring ecological statistics. Proceedings Eighth IEEE International Conference on Computer Vision. ICCV 2001, 2, 416–423 vol.2.

McManus, J. N. J., Li, W., & Gilbert, C. D. (2011). Adaptive shape processing in primary visual cortex. Proceedings of the National Academy of Sciences, 108(24), 9739–9746.

Oleskiw, T. D., Lieber, J. D., Simoncelli, E. P., & Movshon, J. A. (2025). Foundations of visual form selectivity in macaque areas v1 and v2. bioRxiv, 2024.03.04.583307.

Pettersen, K. H., Devor, A., Ulbert, I., Dale, A. M., & Einevoll, G. T. (2006). Current-source density estimation based on inversion of electrostatic forward solution: Effects of finite extent of neuronal activity and conductivity discontinuities. Journal of Neuroscience Methods, 154(1-2), 116–33.

Ponce, C. R., Hartmann, T. S., & Livingstone, M. S. (2017). End-stopping predicts curvature tuning along the ventral stream. The Journal of Neuroscience, 37(3), 648–659.

Pu, S., Dang, W., Qi, X.-L., & Constantinidis, C. (2024). Prefrontal neuronal dynamics in the absence of task execution. Nature Communications, 15(1), 6694.

Quiroga, R. Q., Reddy, L., Kreiman, G., Koch, C., & Fried, I. (2005). Invariant visual representation by single neurons in the human brain. Nature, 435(7045), 1102–1107.

Roe, A. W., & Ts’o, D. Y. (2015). Specificity of v1–v2 orientation networks in the primate visual cortex. Cortex, 72, 168–178.

Rule, M. E., Loback, A. R., Raman, D. V., Driscoll, L. N., Harvey, C. D., & O’Leary, T. (2020). Stable task information from an unstable neural population. eLife, 9, e51121.

Rule, M. E., & O’Leary, T. (2022). Self-healing codes: How stable neural populations can track continually reconfiguring neural representations. Proceedings of the National Academy of Sciences, 119(7), e2106692119.

Rust, N. C., & DiCarlo, J. J. (2010). Selectivity and tolerance (“invariance”) both increase as visual information propagates from cortical area v4 to it. The Journal of Neuroscience, 30(39), 12978–12995.

Sayim, B., & Cavanagh, P. (2011). What line drawings reveal about the visual brain. Frontiers in Human Neuroscience, 5.

Schrimpf, M., Kubilius, J., Hong, H., Majaj, N. J., Rajalingham, R., Issa, E. B., Kar, K., Bashivan, P., Prescott-Roy, J., Schmidt, K., Yamins, D. L. K., & DiCarlo, J. J. (2018). Brain-score: Which artificial neural network for object recognition is most brain-like? bioRxiv, 407007.

Shams, L., & von der Malsburg, C. (2002). The role of complex cells in object recognition. Vision Research, 42(22), 2547–2554.

Simonyan, K., & Zisserman, A. (2015). Very deep convolutional networks for large-scale image recognition. International Conference on Learning Representations.

Skottun, B. C., Bradley, A., Sclar, G., Ohzawa, I., & Freeman, R. D. (1987). The effects of contrast on visual orientation and spatial frequency discrimination: A comparison of single cells and behavior. J Neurophysiol, 57(3), 773–86.

Smith, M. A., Kelly, R. C., & Lee, T. S. (2007). Dynamics of response to perceptual pop-out stimuli in macaque v1. Journal of Neurophysiology, 98(6), 3436–3449.

Spelke, E. S. (2022). What babies know: Core knowledge and composition, vol. 1. Oxford University Press.

Tacchetti, A., Isik, L., & Poggio, T. A. (2018). Invariant recognition shapes neural representations of visual input. Annual Review of Vision Science, 4(Volume 4, 2018), 403–422.

Tanaka, M. (2007). Recognition of pictorial representations by chimpanzees (pan troglodytes). Animal Cognition, 10(2), 169–179.

Tang, S., Lee, T. S., Li, M., Zhang, Y., Xu, Y., Liu, F., Teo, B., & Jiang, H. (2018). Complex pattern selectivity in macaque primary visual cortex revealed by large-scale two-photon imaging. Curr Biol, 28(1), 38–48.e3.

Troyer, T. W., Krukowski, A. E., Priebe, N. J., & Miller, K. D. (1998). Contrast-invariant orientation tuning in cat visual cortex: Thalamocortical input tuning and correlation-based intracortical connectivity. J Neurosci, 18(15), 5908–27.

Vanni, S., Hokkanen, H., Werner, F., & Angelucci, A. (2020). Anatomy and physiology of macaque visual cortical areas v1, v2, and v5/mt: Bases for biologically realistic models. Cereb Cortex, 30(6), 3483–3517.

Victor, J. D., Mechler, F., Repucci, M. A., Purpura, K. P., & Sharpee, T. (2006). Responses of v1 neurons to two-dimensional hermite functions. J Neurophysiol, 95(1), 379–400.

Vinje, W. E., & Gallant, J. L. (2000). Sparse coding and decorrelation in primary visual cortex during natural vision. Science, 287(5456), 1273–1276.

Wilson, E. B. (1927). Probable inference, the law of succession, and statistical inference. Journal of the American Statistical Association, 22(158), 209–212.

Yan, Y., Rasch, M. J., Chen, M., Xiang, X., Huang, M., Wu, S., & Li, W. (2014). Perceptual training continuously refines neuronal population codes in primary visual cortex. Nat Neurosci, 17(10), 1380–7.

Yonas, A., & Arterberry, M. E. (1994). Infants perceive spatial structure specified by line junctions. Perception, 23, 1427–1435.

Ziemba, C. M., Freeman, J., Movshon, J. A., & Simoncelli, E. P. (2016). Selectivity and tolerance for visual texture in macaque v2. Proceedings of the National Academy of Sciences, 113(22), E3140–E3149.

Ziemba, C. M., Freeman, J., Simoncelli, E. P., & Movshon, J. A. (2018). Contextual modulation of sensitivity to naturalistic image structure in macaque v2. Journal of Neurophysiology, 120(2), 409–420.

Ziemba, C. M., Perez, R. K., Pai, J., Kelly, J. G., Hallum, L. E., Shooner, C., & Movshon, J. A. (2019). Laminar differences in responses to naturalistic texture in macaque v1 and v2. The Journal of Neuroscience, 39(49), 9748–9756.

Zipser, K., Lamme, V. A. F., & Schiller, P. H. (1996). Contextual modulation in primary visual cortex. The Journal of Neuroscience, 16, 7376–7389.

